# Reconstruction of cell lineages and behaviors underlying arthropod limb outgrowth with multi-view light-sheet imaging and tracking

**DOI:** 10.1101/112623

**Authors:** Carsten Wolff, Jean-Yves Tinevez, Tobias Pietzsch, Evangelia Stamataki, Benjamin Harich, Stephan Preibisch, Spencer Shorte, Philipp J. Keller, Pavel Tomancak, Anastasios Pavlopoulos

## Abstract

During development coordinated cell behaviors orchestrate tissue and organ morphogenesis to suit the lifestyle of the organism. We have used here the crustacean *Parhyale hawaiensis* to study the cellular basis of limb development. Transgenic *Parhyale* embryos with fluorescently labeled nuclei were imaged at high spatiotemporal resolution with multi-view light-sheet fluorescence microscopy over several days of embryogenesis spanning appendage morphogenesis from early specification up to late differentiation stages. Cell tracking with a new tool called Massive Multi-view Tracker (MaMuT) enabled the reconstruction of the complete cell lineage of an outgrowing thoracic limb with single-cell resolution. *In silico* clonal analyses suggested that the limb primordium becomes subdivided from an early stage first into anterior-posterior and then into dorsal-ventral compartments whose boundaries intersect at the distal tip of the growing limb. Limb bud formation is associated with the spatial modulation of cell proliferation, while limb elongation is also driven by the preferential orientation of division of epidermal cells along the proximal-distal axis of growth. Cellular reconstructions were predictive of the expression patterns of limb development genes including the Decapentaplegic (Dpp) morphogen.

**HIGHLIGHTS:**

- Multi-view light-sheet microscopy of crustacean embryos from species *Parhyale hawaiensis* are ideal for cellular-level analysis of organ morphogenesis.
- Lineages of 3-dimensional organs were reconstructed at single-cell resolution with the Fiji/ImageJ plugin Massive Multi-view Tracker.
- The *Parhyale* limb primordium undergoes early lineage restrictions associated with particular cell behaviors and patterns of gene expression.
- Differential rates of cell proliferation and oriented cell divisions guide appendage proximal-distal outgrowth.

## INTRODUCTION

Developmental morphogenesis is the formation of tissues and organs with particular sizes and shapes to suit the lifestyle of multicellular organisms (Lecuit and Le Goff, 2007). Morphogenesis is driven by patterned cell activities. Therefore, only detailed descriptions of cell lineages and cell behaviors can provide a firm ground for any morphogenetic analysis (Buckingham and Meilhac, 2011). Furthermore, morphogenesis requires capturing and integrating information across multiple levels of biological organization: from the subcellular level, to cell behaviors, to emerging biological form (Keller, 2013; Liu and Keller, 2016).

In this study, we have focused on the crustacean amphipod *Parhyale hawaiensis* that satisfies a number of appealing biological and technical requirements as an experimental model system to study appendage (limb) morphogenesis at single-cell resolution from early specification until late differentiation stages (Stamataki and Pavlopoulos, 2016). First, *Parhyale* is a direct developer; most aspects of the adult body plan, including appendages, are specified during the 10 days of embryogenesis when imaging is readily possible (Browne et al., 2005). Second, *Parhyale* exhibits a striking morphological gradation along the main body axis; each embryo develops a variety of specialized appendages along the anterior-posterior axis (e.g. antennae, mouthparts, limbs for defense, walking, swimming etc.) that differ in size, shape and pattern (Martin et al., 2016; Pavlopoulos et al., 2009; Wolff and Scholtz, 2008). Third, *Parhyale* eggs have the appropriate size and optical properties for microscopic live imaging of constituent cells with very high spatial and temporal resolution; embryos are about 500 *μ*m long, transparent with low autofluorescence and light scattering. Finally, an increasing number of functional genetic approaches, embryological treatments, genomic and transcriptomic resources allow diverse experimental manipulations of *Parhyale* embryos (Kao et al., 2016).

Transient fluorescent labeling by injecting early stage *Parhyale* embryos with mRNAs encoding fluorescent markers coupled with standard epifluorescence or confocal microscopy has been widely adopted to study early processes like gastrulation and germband formation (Hannibal et al., 2012). However, to achieve a comprehensive coverage of appendage formation, *Parhyale* embryos need to be imaged ideally from multiple angular viewpoints and continuously from day 3 up to at least day 8 of embryogenesis (Browne et al., 2005). In this article, we demonstrate that transgenic *Parhyale* embryos with fluorescently labeled nuclei can be imaged routinely for several consecutive days using Light-sheet Fluorescence Microscopy (LSFM). LSFM is an ideal technology for studying how cells form tissues and organs in intact developing embryos (Huisken et al., 2004; Keller et al., 2008; Truong et al., 2011). It enables biologists to capture fast and dynamic developmental processes at very high spatiotemporal resolution, over long periods of time, and with minimal bleaching and photo-damage (Khairy and Keller, 2011; Schmied et al., 2014; Weber et al., 2014). In addition, samples can be optically sectioned from multiple angles (multi-view LSFM) that can be combined computationally to reconstruct the entire specimen with a more isotropic resolution (Chhetri et al., 2015; Krzic et al., 2012; Swoger et al., 2007; Tomer et al., 2012; Wu et al., 2013).

Although the amount and type of data generated by multi-view LSFM raise several challenges for image processing and analysis, most of them have been efficiently addressed. Software solutions exist for registration of acquired views (z-stacks) in each time-point (Preibisch et al., 2010), for fusion of views into a single output z-stack with nearly isotropic resolution (Preibisch et al., 2014), and for rendering and three dimensional reconstruction of the entire imaged volume (Pietzsch et al., 2015). These processes can be repeated automatically using high-performance computing resources for several hundred or thousands of time-points to generate a four-dimensional representation (three spatial dimensions + temporal dimension) of the embryo as it develops over time (Amat et al., 2015; Schmied et al., 2014). Automated approaches for cell segmentation and tracking have also been developed (Amat et al., 2014), however they do not yet reach the precision required for unsupervised extraction of cell lineages. To address this issue, we describe here the Massive Multi-view Tracker (MaMuT) software that allows the visualization, annotation, and accurate lineage reconstruction of large multi-dimensional microscopy data.

We quantitatively followed cell dynamics with the MaMuT software on the *Parhyale* multi-view LSFM acquisitions in order to understand the cellular basis of arthropod appendage morphogenesis. As revealed by lineage tracing experiments in the leading arthropod model *Drosophila melanogaster,* the leg and wing primordia become progressively subdivided into distinct cell populations (called compartments when lineage-restricted), first along the anterior-posterior (AP) axis during their specification in the early embryo, and later along the dorsal-ventral (DV) axis during larval stages (Dahmann et al., 2011; Garcia-Bellido et al., 1973; Steiner, 1976). These lineage-dependent or independent subdivisions of the tissue acquire distinct cell fates or identities driven by domain-specific expression of patterning genes (called selector genes if lineally inherited), as well as by the localized expression of signaling molecules at compartment boundaries (“organizers”) that control patterning and growth of the developing organs (Garcia-Bellido, 1975; Lawrence and Struhl, 1996; Mann and Carroll, 2002; Restrepo et al., 2014). Besides these regionalization mechanisms, oriented cell divisions have been implicated as a general mechanism in shaping the *Drosophila* wing and other growing organs in diverse developmental systems (Baena-Lopez et al., 2005). Other cellular activities like patterned cell proliferation and cell rearrangement could also play a role in the formation of limb buds and their elongation along the proximal-distal axis, but these processes have been difficult to study in existing arthropod models. By acquiring the lineage information at single-cell resolution, as well as the positional coordinates and their change over time for all constituent cells in a developing limb, we identified the lineage restrictions and morphogenetic cellular behaviors operating during *Parhyale* limb bud formation and elongation, and compared these to the *Drosophila* and other arthropod paradigms. Finally, we validated our reconstructions by studying at cellular resolution the expression of Decapentaplegic (Dpp) signaling components implicated in limb patterning and growth.

## RESULTS

### Imaging *Parhyale* embryogenesis with multi-view LSFM

Segment formation and maturation in *Parhyale* occurs sequentially in anterior-to-posterior progression (Browne et al., 2005). In our LSFM recordings, we were particularly interested in capturing appendage development in the anterior thorax of *Parhyale* embryos. In these segments, appendages were specified at about 3.5 days after egg-lay (AEL) at 25°C. Over the next 4 days, appendage buds bulged out ventrally, elongated along their proximal-distal axis and became progressively segmented until they acquired their definite morphology at around 8 days AEL (Figure 1).

**Figure 1.**
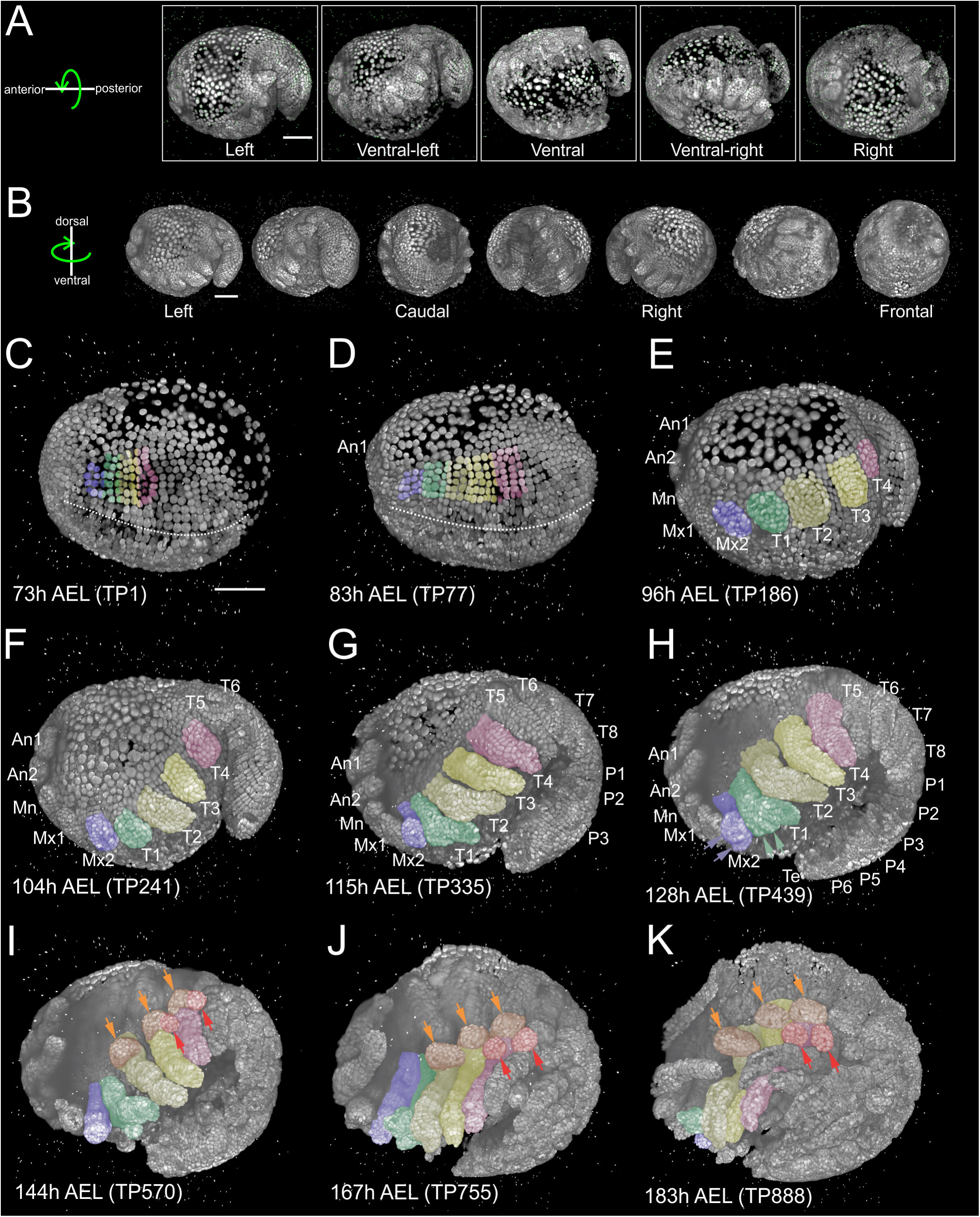
Reconstruction of *Parhyale* embryogenesis from multi-view LSFM. (A) Transgenic *Parhyale* embryos with H2B-mRFPruby labeled nuclei were mounted in agarose with scattered red fluorescent beads (green dots) as fiducial markers for multi-view reconstruction. Each embryo was imaged from the indicated 5 views with a 45° rotation around the anterior-posterior body axis between neighboring views. Each panel shows a 3D rendering of the corresponding view with anterior towards the left. (B) Input views were registered and fused computationally into a single output image rendered here in different positions around the dorsal-ventral axis. (C-K) Each panel shows a 3D rendering of the fused volume at the indicated developmental stage in hours (h) after egg-lay (AEL) and the corresponding time-point (TP) of the recording. Abbreviations: first antenna (An1), second antenna (An2), mandible (Mn), maxilla 1 (Mx1), maxilla 2 (Mx2), thoracic appendages 1 to 8 (T1-T8), pleonic (abdominal) appendages 1 to 6 (P1-P6) and telson (Te) at the posterior terminus of the body. Color masks are used to indicate the cells contributing to the developing Mx2 (in blue), T1 maxilliped (in green), T2 and T3 gnathopods (in light and dark yellow, respectively) and T4 limb (in magenta). (C) Embryo at mid-germband stage (S13 according to (Browne et al., 2005)). The ventral midline is denoted with the dotted line. (D) Embryo at stage S15. Germband has extended to the posterior egg pole and the first antennal bud is visible anteriorly. (E) Embryo at stage S18 with posterior ventral flexure. Head and thoracic appendages have bulged out up to T4. (F) Embryo at stage S19 with prominent head and thoracic appendage buds up to T6. (G) Embryo at stage S20 continues elongating ventrally and anteriorly. Appendage buds are also visible in the pleon up to P3. (H) Embryo at stage S21. Segmentation is complete and all appendages have formed. The Mx2 has split into two branches (blue arrowheads) and the T1 maxilliped has developed two proximal ventral outgrowths (green arrowheads). (I) Embryo at stage S22, (J) stage S23, and (K) stage S24 showing different stages of appendage segmentation. Dorsal outgrowths at the base of thoracic appendages, namely coxal plates (orange arrowheads) and gills (red arrowheads), are indicated in T2, T3 and T4. In all panels anterior is towards the left and dorsal towards the top. Scale bars are 100 *μ*m.

We generated transgenic *Parhyale* embryos with fluorescently labeled nuclei as described in Materials & Methods. Individual 3-day old embryos (stage S13) were mounted for LSFM in a column of low melting agarose with scattered red fluorescent beads. Embryos were imaged on a Zeiss Lightsheet Z.1 microscope that offers dual-sided illumination with single-sided detection. Several parameters (detailed in Materials & Methods) were optimized to cover all stages of *Parhyale* appendage development over 4-5 days of embryogenesis at single-cell resolution with adequate temporal sampling and with minimal photo-bleaching and damaging. A typical 4 to 5-day long *Parhyale* embryo recording was composed of more than 1 million images resulting in >7 TB datasets.

The slow tempo of *Parhyale* development enabled LSFM imaging of the entire embryo from multiple highly overlapping views (every 45°, Figure 1A) without noticeable displacement of cell nuclei between all the views acquired in each time-point. Development of the entire embryo was reconstructed from the raw input views according to the following steps (detailed in Materials & Methods) using open-source software available as plug-ins in the Fiji (Fiji Is Just ImageJ) biological image analysis platform (Schindelin et al., 2012): 1) image file preprocessing, 2) bead-based spatial registration of views within one time-point, 3) fusion by multi-view deconvolution, 4) bead-based temporal registration across time-points, 5) computation of temporally registered fused volumes, and 6) 4D rendering of the spatiotemporally registered fused data (Preibisch et al., 2014; Preibisch et al., 2010; Schmied et al., 2014). This processing resulted in almost isotropic resolution of fused volumes and was used for visualization of *Parhyale* embryo development with cellular resolution (Figure 1B).

### Following appendage morphogenesis in 4D reconstructed Parhyale embryos

Appendage morphogenesis in arthropods involves patterning, growth and differentiation of ectodermal cells organized in an epithelial monolayer that will give rise to the appendage epidermis and will secrete the cuticle. Our study focused on the dynamics of the *Parhyale* ectoderm from the stages of appendage specification (day 3 AEL) up to differentiation (day 8 AEL). During germband formation, the ectoderm contributing to the posterior head and the trunk (known as post-naupliar germband) became organized in a stereotyped grid-like pattern with a very precise arrangement of rows and columns of cells (Figures 1C-1D and Figures 2A-2A’’) (Browne et al., 2005; Dohle et al., 2004). Each row of cells corresponded to one parasegment, which is the unit of early metameric organization in *Parhyale* embryos, like in *Drosophila* and other arthropods (Hejnol and Scholtz, 2004; Scholtz et al., 1994). Two rounds of longitudinally-oriented cell divisions in each formed parasegmental row (Figures 2B-2D’), together with the progressive addition of new parasegments at the posterior end, led to embryo axial elongation (Figures 1D-1H).

**Figure 2.**
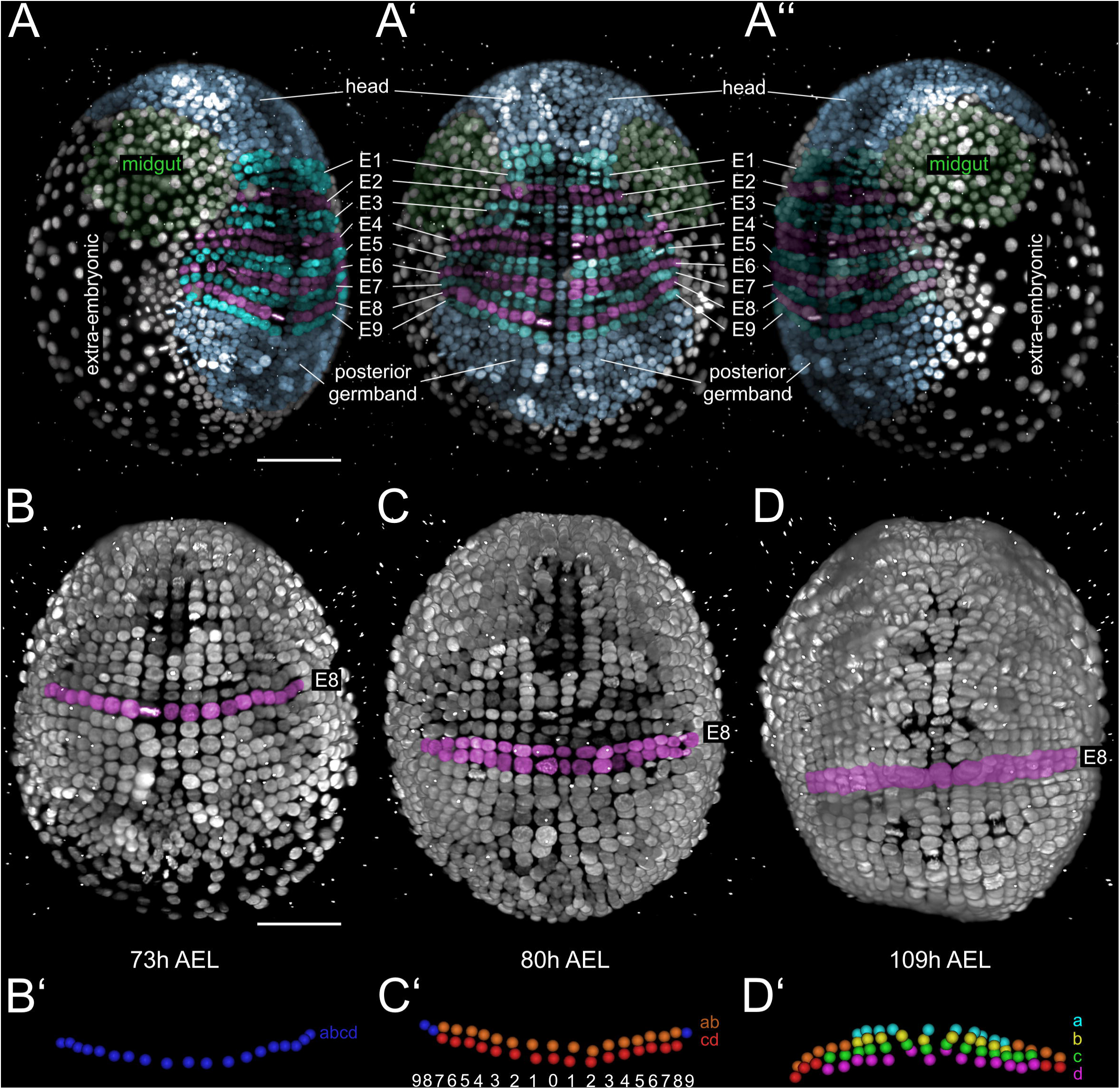
Grid architecture of the *Parhyale* germband. (A-A’’) 3D rendering of *Parhyale* embryo at the growing germband stage: (A) Right side, (A’) ventral side, and (A’’) left side. Color masks indicate the anterior head region (blue), the bilaterally symmetric midgut precursors (green), the orderly arranged parasegments E1 to E10 (in alternating cyan and magenta), the posterior end of the germband (blue) with ongoing organization of cells into new rows, and the extraembryonic tissue (white). (B-D) Ventral views of elongating germband at the indicated hours (h) after egg-lay (AEL). Ectodermal cells making up the E8 parasegment are shown in magenta. (B’) Schematic of tracked E8 abcd cells (blue) at the 1-row-parasegment, (C’) anterior ab cells (orange) and posterior cd cells (red) after the first longitudinally-oriented division at the 2-row-parasegment, and (D’) a cells (cyan), b cells (yellow), c cells (green) and d cells (magenta) after the second longitudinally-oriented division at the 4-row-parasegment. Both mitotic waves proceed in medial to lateral direction. The resulting daughter cells sort in clearly defined columns. In each hemisegment, ascending numbers indicate column position relative to the ventral midline (column 0).

Subsequent divisions of ectodermal cells had a more complex pattern disrupting the regularity of the grid and contributing to the transition from parasegmental to segmental body organization with the formation of segmental borders and the evagination of paired appendages in each segment. Appendage buds appeared successively from the head region backwards (Figures 1D-1H) and started lengthening (Figures 1F-1K) and differentiating along their proximal-distal axis (Figures 1G-1K). The gnathal appendages (mandible, maxilla 1, maxilla 2) grew relatively little in size, remained unsegmented and bifurcated heralding the development of gnathal palps (Figure 1H). The most conspicuous appendages of the embryo were the two antennae and the thoracic appendages. These appendages elongated considerably along their proximal-distal axis and became progressively subdivided into distinct segments (Figures 1F-1K and Figure 6). At the end of the imaging period, morphogenesis appeared nearly complete. All thoracic appendages (T1 to T8) were composed of 7 constituent segments that had distinct pattern, size and shape to serve their specialized function: T1 maxillipeds for feeding, T2 and T3 gnathopods for grasping and T4 to T8 legs for locomotion. Besides elongation and segmentation, thoracic appendages also developed a number of different proximal outgrowths. The maxillipeds developed two ventral outgrowths, called endites (Figure 1H), while the more posterior thoracic appendages developed dorsal outgrowths, called epipodites, serving as protective coxal plates and respiratory gills (Figures 1I-1K).

Thus, multi-view LSFM imaging captures the entire gamut of differential appendage morphogenetic events along the body axis of the *Parhyale* embryo in a single time-lapse experiment.

### MaMuT: a platform for cell tracking in multi-view and multi-terabyte datasets

In order to examine the cellular basis of morphogenesis, we developed a novel Fiji software application to extract cell lineages from multi-view and multi-terabyte datasets. This tool was dubbed MaMuT for Massive Multi-view Tracker and is a hybrid and extension of two existing Fiji plugins: the BigDataViewer visualization engine (Pietzsch et al., 2015); http://imagej.net/BigDataViewer) and the TrackMate annotation engine (Tinevez et al., 2016); http://imagej.net/TrackMate). MaMuT is an interactive, user-friendly tool for visualization, annotation, object tracking and lineage reconstruction of large multi-dimensional microscopy data. Within the Fiji ecosystem, MaMuT is tightly integrated with the plugins for multi-view LSFM data processing (Figure 3A). Documentation, training datasets and tutorials on MaMuT can be found at the dedicated link http://imagej.net/MaMuT.

**Figure 3.**
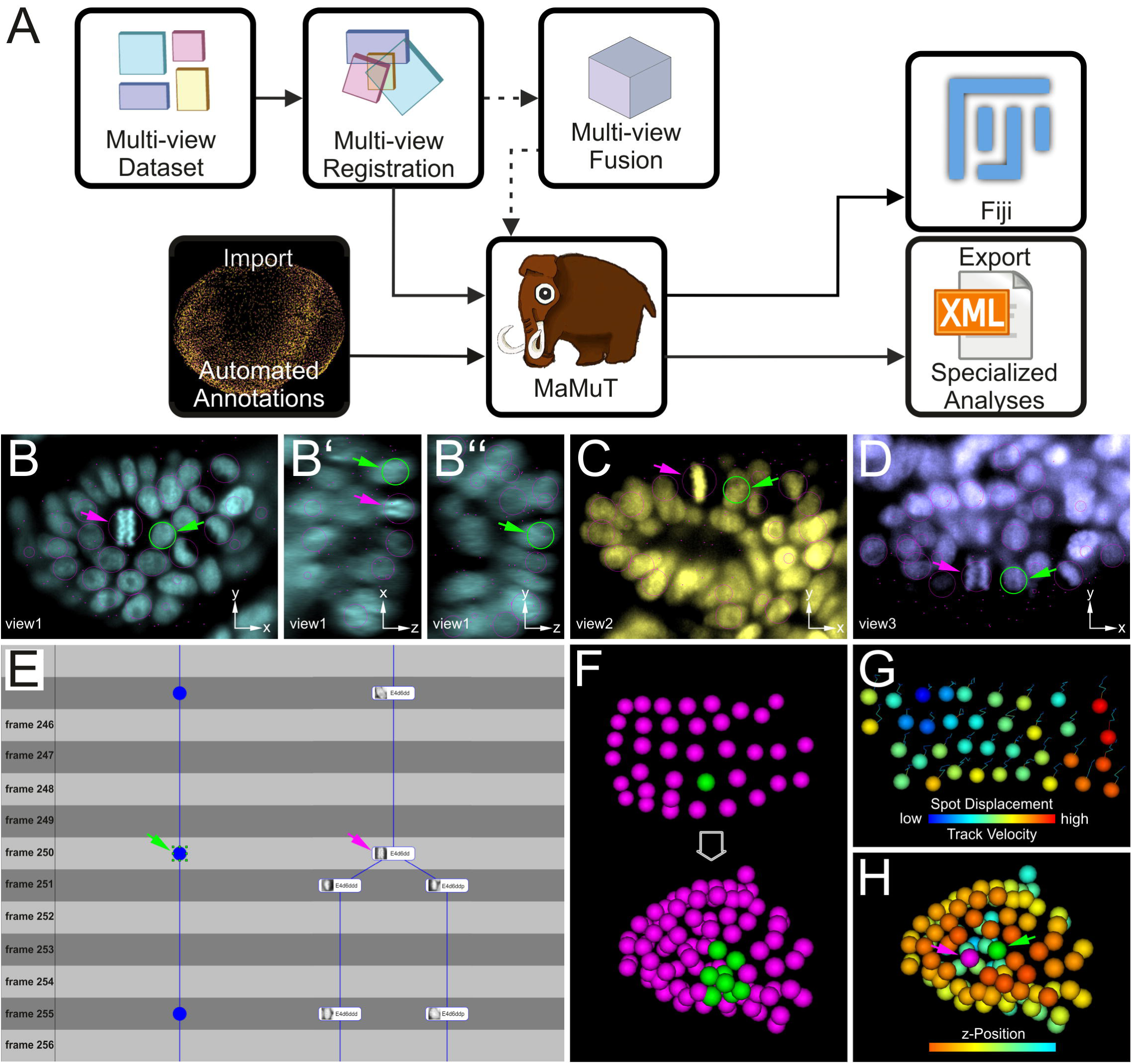
Lineage reconstruction and cell tracking with MaMuT. (A) Workflow for image data analysis with the Massive Multi-view Tracker (MaMuT) Fiji plugin. The raw views/z-stacks (colored boxes in Multi-view Dataset) are registered (overlapping boxes in Multi-view Registration) and, optionally, fused into a single volume (large cube in Multi-view Fusion). The raw (and/or fused) image data together with the registration parameters are imported into MaMuT (represented with mammoth logo). In its simplest implementation, all lineaging, tracking and visual interpretation of the data can be done with MaMuT in the Fiji workspace. In more advanced implementations, the image and registration data are complemented with automated segmentation and tracking annotations computed separately (Import of Automated Annotations pictured as point cloud of tracked cells). The reconstructed positional, temporal and lineage information can be exported from MaMuT in an xml file and imported into other platforms for more specialized analyses. (B-D) Viewer windows displaying the raw *Parhyale* image data and annotations. All selected nuclei are marked with magenta circles (in view) or dots (out of view). The currently selected nucleus is marked in green in all Viewers: (B) xy plane, (B’) xz plane, and (B’’) yz plane of the first available ventral-lateral view in cyan; (C) xy plane of the second available ventral view in yellow; (D) xy plane of the third available lateral view in blue. (E) Close up of the TrackScheme lineage browser and editor where tracks are arranged horizontally and time-points are arranged vertically. Tracked objects can be displayed simply as spots (left track) or with extra information like their names and thumbnails (right track). Tracks are displayed as vertical links. The TrackScheme is synced with the Viewer windows, thus the selected nucleus is also highlighted here in green at the indicated time-point (frame). Note that objects can be tracked between consecutive time-points (e.g. in right track) or in larger steps (e.g. in left track). (F-H) Animations of tracked objects depicted as spheres in the 3D Viewer window: (F) Digital clone of a nucleus (shown in green) tracked from the grid stage to the limb bud stage. All other tracked nuclei are shown in magenta. (G) Spots and tracks can be color-coded independently according to various numerical parameters extracted from the data, like displacement (spot colors), velocity (track colors), mean cell division time (not shown) and others. (H) Tracked nuclei at the limb bud stage mapped out in different colors based on z-position (dorsal-ventral position in this case). In panels B-H, the selected nucleus and the neighboring dividing nucleus are indicated with green and magenta arrowheads, respectively.

MaMuT can handle multiple data sources but was developed primarily to enable the analysis of large and complex LSFM image datasets (Figure 3 and Figure S1). Its unique feature is the ability to annotate an image volume synergistically from all available input views without the need to fuse these into a single volume. All raw views were first registered and then imported into MaMuT (Figure 3A). The software allowed users to open the raw image data in as many *Viewer* windows as required and visualize each z-stack in any desired orientation, scale and time-point (Figures 3B-3D). Most importantly, all *Viewer* windows were synced based on the calculated registration parameters and shared a common physical coordinate system. That is, upon selecting an object of interest, like a cell or nucleus (“spot”) in one *Viewer,* the same spot was identified and displayed in all other windows, and its x, y, z position was mapped onto this common physical space (Figures 3B-3D). This unique functionality of MaMuT allowed us to identify and track all constituent cells in a developing appendage continuously from the early germband stages until the later stages of 3D organ outgrowth (Figures 4 to 6), when the information from multiple views was required to fully reconstruct the appendage with single-cell resolution. Selected nuclei were tracked over time in user-defined time intervals, and the reconstructed trajectories and lineages were also displayed in two additional synced windows, the *TrackScheme* and *3D Viewer.* The *TrackScheme* lineage browser and editor displayed the reconstructed cell lineage tree with tracked nuclei represented as nodes connected by edges over time and cell divisions depicted as split branches in the tree (Figure 3E). The *3D Viewer* window displayed interactive animations of the spots depicted as spheres and their tracks over time (Figure 3F-3H). The spots and the tracks in the *Viewer*, *TrackScheme* and *3D Viewer* windows could be color-coded by lineage, position and other numerical features to assist visual analysis and interpretation of the data (Figures 3F-3H and Figures 4 to 7). In addition, all these windows were synced to simultaneously highlight selected spots of interest at the selected time-point, greatly facilitating the cell lineaging process (Figures 3B-3H).

**Figure 4.**
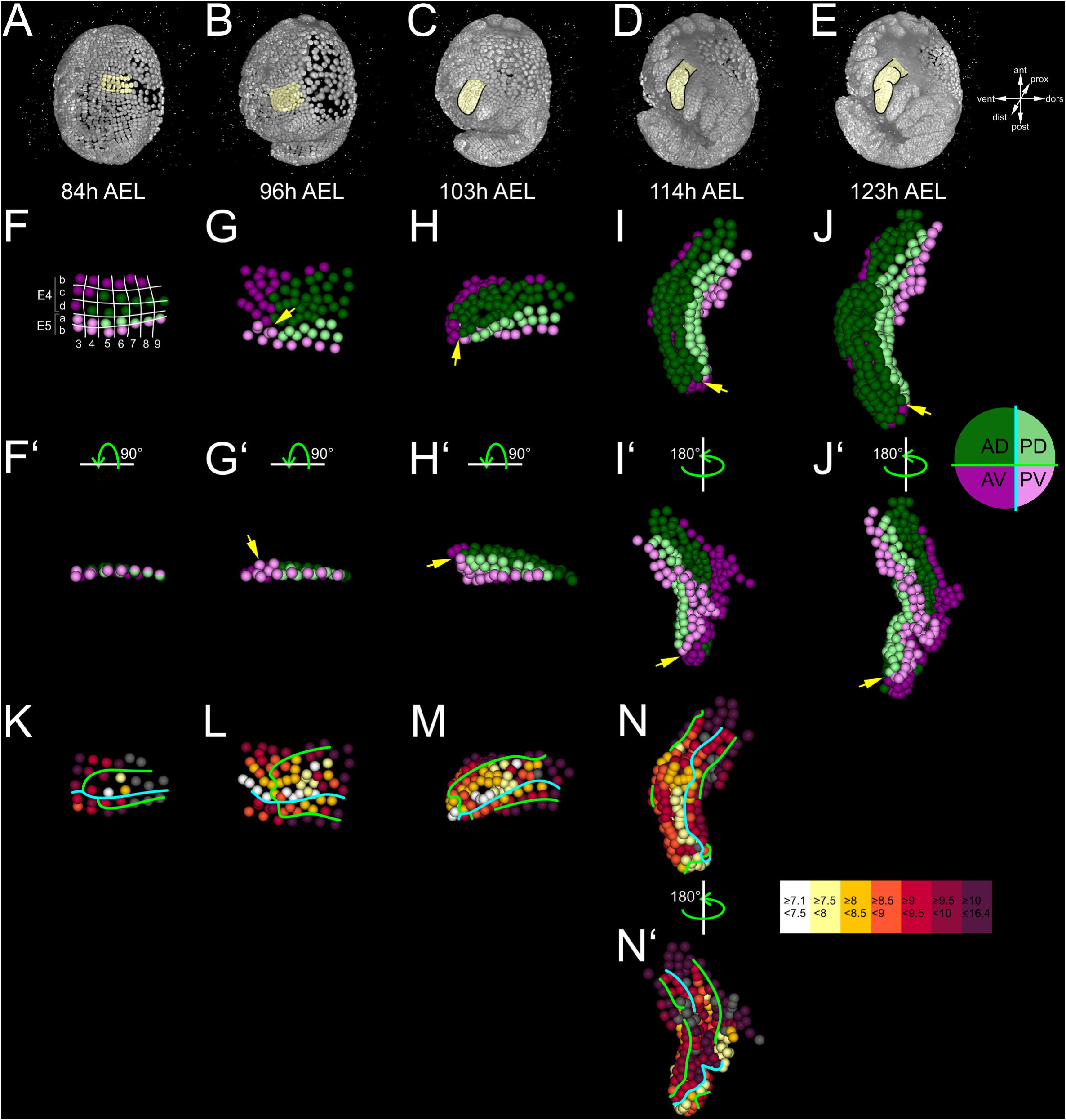
Cellular architecture and dynamics in the *Parhyale* thoracic limb. (A-E) Lateral views of a *Parhyale* embryo reconstructed from a multi-view LSFM acquisition at the indicated developmental stages shown in hours (h) after egg-lay (AEL). The yellow mask indicates the second thoracic appendage (T2) on the left side that was reconstructed at a single-cell resolution. (F-J’) Tracked cells contributing to the outgrowing T2 limb color-coded by their compartmental identity: anterior-dorsal (AD; dark green), anterior-ventral (AV; dark magenta), posterior-dorsal (PD; light green), and posterior-ventral (PV; light magenta). (F) Ventral view of the limb primordium at the 4-row-parasegment stage at 84 h AEL. The T2 primordium is made up by cells from the E4 and E5 parasegments. Horizontal lines separate rows a to d along the AP axis, and vertical lines separate columns 3 to 9 along the DV axis. (F’) Posterior view, rotated 90° relative to F. (G) Ventral view of the limb primordium during early eversion at 96 h AEL. (G’) Posterior view, rotated 90° relative to G. The cells close to the intersection of the four compartments (yellow arrow) are the first to rise above the level of the epithelium. (H-J) Dorsal views of (H) limb bud at 103 h AEL, (I) initial limb elongation at 114 h AEL and (J) later elongation phase at 123 h AEL. (H’) Posterior view, rotated 90° relative to H, and (I’-J’) ventral views, rotated 180° relative to I-J. The intersection of the AP and DV compartment boundaries (yellow arrows) is located at the tip of the limb. (K-N’) Same stages and views as in panels F-I and I’ color-coded by the average cell cycle length of each cell according to the scale shown on the right. The AP and DV compartment boundaries are indicated by the cyan and green line, respectively. Gray cells indicate cells for which measurements are not applicable. (K) Some central c and d cells start dividing faster at the 4-row-parasegment stage. (L) During early eversion, the middle cells divide faster compared to peripheral cells. (M) At limb bud stage, higher proliferation rates are detected at the tip and in the anterior-dorsal compartment. (N) Dorsal and (N’) ventral view of elongating limb. Cells at the tip of the limb and anterior cells abutting the AP compartment boundary divide the fastest.

Although, all lineage reconstructions presented in this article were generated manually, the latest MaMuT architecture also offers two functionalities for automated tracking: i) a semi-automated option where individual nuclei can be selected by the user and segmented and tracked computationally over time, and ii) the option to import into MaMuT fully automated annotations generated by the Tracking with Gaussian Mixture Models (TGMM) software (Amat et al., 2014), which is one of the most accurate and computationally efficient methods for segmentation and tracking of fluorescently labeled nuclei (Figure 3A). Therefore, MaMuT is a versatile platform that can be used either for fully manual or semi-automated tracking of selected populations of objects of interest, or for visualization and editing of fully automated computational predictions for systems-level lineage reconstructions.

### Single-cell lineage reconstruction of a Parhyale thoracic limb

We next deployed the manual version of MaMuT to extract the developmental lineage tree of a *Parhyale* thoracic limb. By convention, ectodermal rows in the *Parhyale* germband are identified from anterior to posterior by ascending numbers E0, E1, E2 etc. (Figures 2A-2A’’). Each parasegmental row of cells, labeled abcd (Figures 2B-2B’), undergoes two rounds of divisions in anterior-posterior direction to first generate two rows, labeled ab anteriorly and cd posteriorly (Figures 2C-2C’), and then four rows of cells labeled a, b, c and d from anterior to posterior (Figures 2D-2D’). Columns of ectodermal cells are identified by ascending index numbers with 0 denoting the ventral midline and 1, 2, 3 etc. denoting the more lateral columns with increasing distance from midline (Figure 2C’).

In accordance with previous studies in malacostracan crustaceans and various other arthropods, our reconstructions demonstrated that each *Parhyale* thoracic limb consisted of cells from two neighboring parasegments (Browne et al., 2005; Dohle et al., 2004; Scholtz et al., 1994; Wolff and Scholtz, 2008). The T2 limb that we analyzed in-depth (Figure 4A-4E) developed from rows b, c and d of the E4 parasegment and from rows a and b of the following E5 parasegment (Figure 4F and Figure S2). Cells that arose from rows c, d and a occupied the entire length of the limb proximal-distal axis and the body wall part of the T2 segment (Figure S2). Descendent cells from rows b contributed only to the proximal limb and intersegmental territories (Figure S2). Cells in medial columns 1 and 2 gave rise to the nervous system and sternites and were not considered further in this study. The more lateral columns 3 to 9 contributed the epidermal cells forming the limb (Figure S3).

We fully tracked 34 cells constituting the T2 limb primordium over 50 hours of *Parhyale* embryogenesis giving rise to a total of 361 epidermal cells. We started tracking each of these 34 cells as they divided longitudinally from the 2-row to the 4-row parasegment and monitored them continuously during the subsequent rounds of divisions (referred to as differential divisions, DDs). The number of DDs observed during these 50 hours (400 time-points) varied dramatically between tracked cells from just one DD in the slowest dividing lateral cells of the T2 primordium (cells E4b8, E5a9/b9) to five DDs in the fastest dividing central cells (cells E4c3-c6 and E4d3-d6). Although the clonal composition of crustacean appendages had been described previously using single-cell injections with lypophilic dyes (Wolff and Scholtz, 2008), the reconstruction presented here is the most comprehensive lineage tree for any developing arthropod limb published to date (Figure S4).

### Early lineage restrictions along the anterior-posterior and dorsal-ventral axes

We first asked whether these complete reconstructions could reveal any lineage-based subdivisions in the developing limb. The AP restriction at the border of neighboring parasegments from the 1-row-parasegment stage onwards has been revealed in *Parhyale* and other malacostracan embryos by classic embryological descriptions, lineage tracing experiments and expression studies for the segment polarity gene *engrailed* that marks the posterior compartment (Browne et al., 2005; Dohle et al., 2004; Hejnol and Scholtz, 2004; Scholtz et al., 1994). In agreement with this AP restriction, during limb specification, outgrowth and elongation there was a straight clonal boundary running between the anterior compartment cells derived from the E4b, c and d rows and the posterior compartment cells derived from the E5a and b rows (Figure 4 and Figure S2).

Having established that the well-known AP boundary was readily observable in our complete lineages, we next sought to identify any subdivision along the dorsal-ventral axis. Compartments were classically discovered by clonal analysis using mitotic recombination. In our reconstructions, we could generate clones digitally from arbitrary cells at different stages of appendage development. We reasoned that we could reveal the timing and position of any heritable DV restriction by piecing together correctly all founder cells of dorsal or ventral identity in a way that the two polyclones (i.e. compartments) would stay separate and form a lasting straight interface between them. This analysis suggested that there is indeed a DV separation that took place at the 4-row-parasegment stage; the DV boundary ran between the E4b and c rows anteriorly, between the E5a and b rows posteriorly, and between cells E4c4-c5, E4d3-d4 and E5a4-a5 medially (Figure 4F). Throughout limb development, the dorsal and ventral cells formed a sharp boundary between themselves extending along the proximal-distal axis (Figures 4I-4J’ and Figure S2). In order to evaluate the reproducibility and stereotypy of the AP and, in particular, the DV separation across *Parhyale* embryos, we analyzed an independently imaged and reconstructed T2 limb from a different embryo (Figure S5). We confirmed that four identical compartments (anterior-dorsal, anterior-ventral, posterior-dorsal and posterior-ventral) could be derived in this independent reconstruction with straight boundaries and no cell mixing between neighboring compartments (compare Figure 4 with Figure S5).

Taken together, these results suggested that *in silico* analyses of comprehensive and accurate lineage reconstructions can provide novel insights into clonal subdivisions and the underlying developmental patterning mechanisms in species where sophisticated genetic dissections are not implemented yet.

### Cellular dynamics underlying limb bud formation and elongation

At the tissue level, T2 limb bud formation entailed the transformation of a flat two-dimensional epithelial sheet into a three-dimensional bulge (Figures 4A-4C). At the cellular level, the first step in this transformation was the rise of few cells above the level of the germband at 96 hours AEL (stage S18; Figures 4G-4G’). These were the cells abutting the intersection of the four compartments. Within the following 3 hours, this initial phase was followed by a large-scale elevation of most cells in the dorsal compartment. As this elevation continued, the medial ventral cells folded and became apposed to the medial dorsal cells forming the convex surface of the limb bud (Figures 4H-4H’). The intersection of the four compartments was at the tip of the limb bud and persisted in this position throughout subsequent elongation (Figures 4H-4J’). From 103 hours AEL onwards (stage S19), a second element started bulging distally of the original limb bud (Figures 4I-I’ and Figures 6I-6O). The limb elongated as a convoluted rather than straight cylinder and acquired progressively an S-shape (Figures 4J-4J’). Our reconstructions suggested that cell movements played little role, at least during the analyzed stages of limb outgrowth. We identified few instances of cells intercalating between one another along the proximal-distal axis during the extension phase slightly contributing to the narrowing and lengthening of the growing limb (Figure S6).

### Quantification of differential cell behaviors during limb formation

We next asked whether our reconstructions could provide insights into the morphodynamics of limb formation at a finer scale. Two cell behaviors implicated in tissue and organ morphogenesis were readily quantifiable in our nuclear trackings, namely the pattern of cell proliferation and the orientation of cell divisions (Baena-Lopez et al., 2005; Boehm et al., 2010). These cell activities have been traditionally inferred in developing tissues from the distribution, size and shape of somatic clones induced by various means (Buckingham and Meilhac, 2011). This approach could be also adapted here by generating *in silico* clones for every single tracked epidermal cell between any developmental stages of interest (Figure S7). Yet, the MaMuT reconstructions enabled us to enrich the lineage information with rigorous quantitative analyses of the rate and orientation of mitotic divisions using the extracted x, y, z, t coordinates for all tracked nuclei.

First, we calculated the cell cycle length (CCL), i.e. the branch length in the reconstructed cell lineages, for every constituent cell in the developing T2 limb (Figures 4K-4N’ and Figure S8). This quantification revealed a striking difference in CCL between cells in the limb primordium during early limb bud formation. The cells in the center of the primordium were dividing faster than their neighboring cells in the periphery of the primordium (average CCL 7.1-8.5 hours for central cells versus 8.5-16.4 hours for peripheral cells). This difference started from early primordium specification at the 4-row-parasegment (Figure 4K), but became most pronounced during the global elevation of the limb bud cells (Figure 4L), suggesting a causal association between spatially controlled cell proliferation and initiation of limb outgrowth (see Discussion). During subsequent elongation stages, a high concentration of fast dividing cells was located at the intersection of the four presumptive compartments, resembling a growth zone at the distal tip of the growing appendage (Figures 4M-4N). Another stripe of faster dividing cells was discovered in the anterior-dorsal cells abutting the AP compartment boundary (Figures 4N-4N’).

Next, we looked for any biases in the orientation of mitotic divisions that could be associated with *Parhyale* limb morphogenesis similar to the *Drosophila* wing paradigm (Baena-Lopez et al., 2005). The AP compartment boundary ran parallel to the proximal-distal axis of limb growth and provided an excellent axis of reference against which we compared the axis of each cell division (Figures 5). As expected, all early divisions in the T2 primordium were perpendicular to the AP compartment boundary (i.e. they were parallel to the AP axis) confirming the strict longitudinal orientation of row divisions (Figure 5F). Cell divisions acquired a more heterogeneous pattern after the 4-row-parasegment stage (Figure 5G). An increasing number of mitotic spindles aligned progressively along the proximal-distal axis during limb bud formation (Figure 5H) and elongation (Figures 5I-5J).

**Figure 5.**
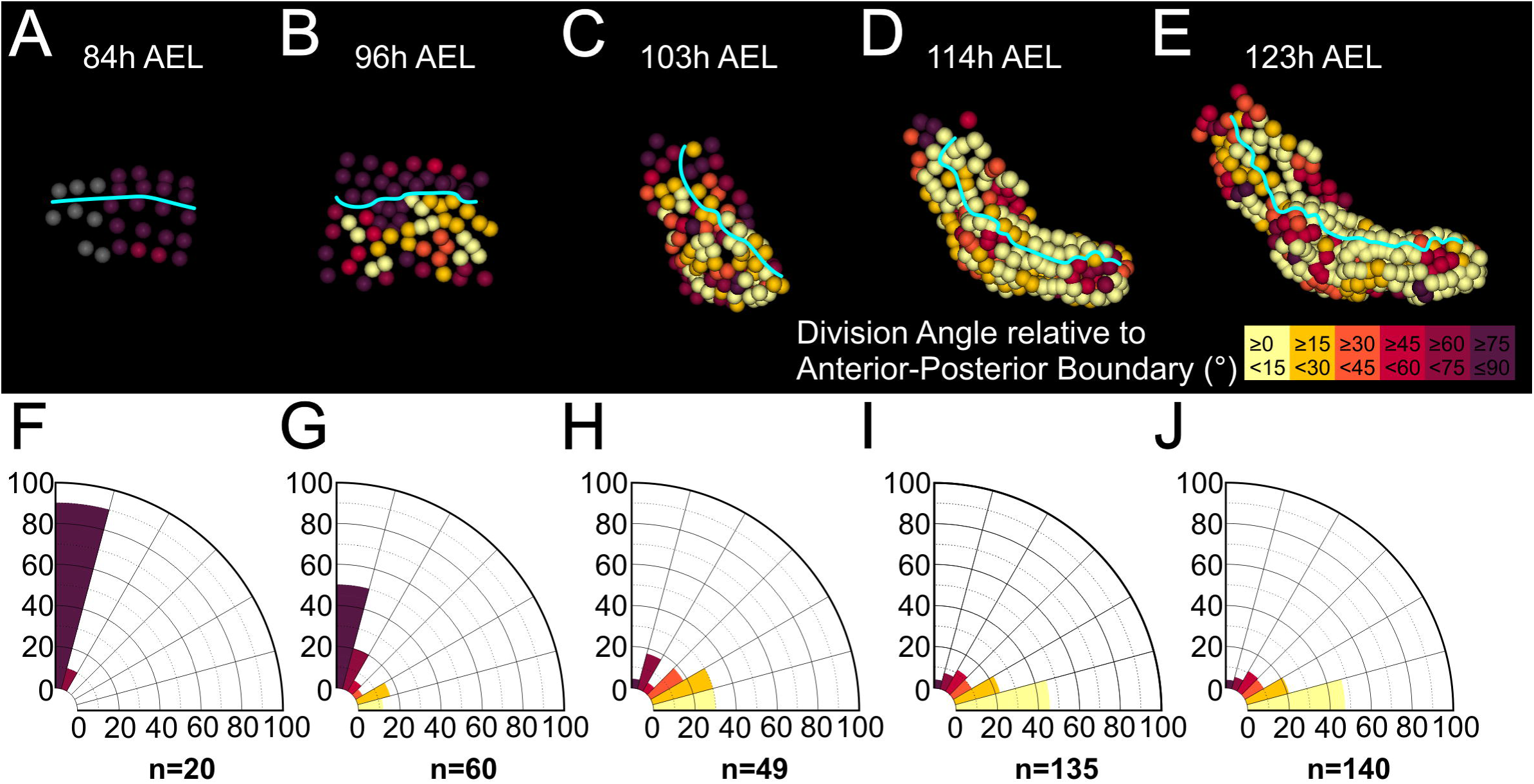
Quantification of orientation of mitotic divisions in the *Parhyale* thoracic limb. (A-E) Tracked cells making up the T2 limb shown at the indicated hours (h) after egg-lay (AEL) and color-coded by the orientation of mitotic divisions relative to the AP compartment boundary (cyan line). The absolute values of the division angle relative to the AP boundary are sorted in 6 bins of 15°. Gray cells in panel A indicate non-divided cells. Note that the AP boundary is an accurate proxy for the proximal-distal axis of limb growth. (F-J) Corresponding rose diagrams with 15° intervals showing the percentage of mitotic events falling in each bin (division angles relative to AP boundary) and color-coded as in A-E. The number of mitotic events (n) between the indicated time-points is shown under the rose diagrams. (A-F) Limb primordium at the 4-row-parasegment stage. Only longitudinally-oriented divisions (i.e. perpendicular to AP boundary) are detected 73 to 85 hours AEL. (B-G) Limb primordium during early eversion. Most cells still divide longitudinally 85 to 96 hours AEL, but an increasing number of dividing cells align parallel to the AP boundary. (C-H) Limb bud stage. More than 59% of cells divide 0°-30° relative to the AP boundary 96 to 103 hours AEL. (D-I) Early and (E-J) later limb elongation phase. The large majority of cells (>68%) divide 0°−30° relative to the AP boundary 103 to 123 hours AEL.

Collectively, the information extracted from our spatiotemporally resolved lineage trees strongly suggested that *Parhyale* limb outgrowth is driven by at least two patterned cell behaviors: the differential rates of cell proliferation and the orderly arrangement of mitotic spindles.

### Cellular basis of the elaboration of the limb proximal-distal axis

We then asked whether our dataset could help resolve the patterning mechanisms operating during appendage segmentation. Appendage segmentation is a gradual process as the elongating proximal-distal axis becomes progressively subdivided into an increasing number of elements (Figure 6) (Rauskolb, 2001). At the tissue level, the forming *Parhyale* T2 limb was made of a single bud up to 103 hours AEL (Figures 6G-6H), while a second element bulged out distally from the original bud at around 114 hours AEL (Figure 6I). Over the next 40 hours, a number of consecutive circumferential constrictions subdivided the limb into the final pattern of 7 segments (Figures 6J-6L) that are called (from proximal to distal) coxa, basis, ischium, merus, carpus, propodus, and dactylus.

**Figure 6.**
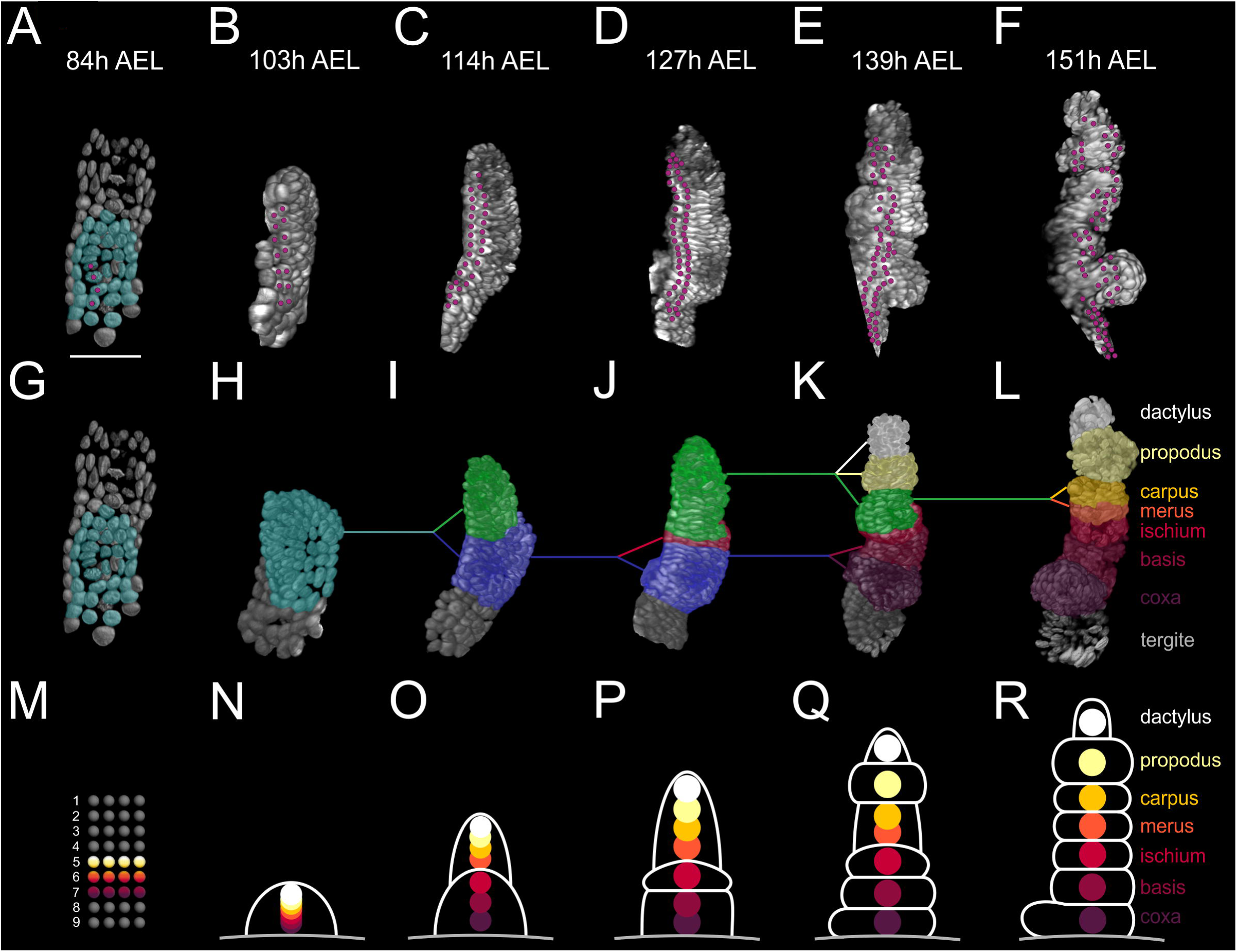
Elaboration of the *Parhyale* limb proximal-distal axis. (A-F) 3D renderings of the T2 limb at the indicated hours (h) after egg-lay (AEL). The cells contributing to the T2 primordium are shown in cyan in panel A. Magenta dots indicate the tracked cells E5a5-a8 and their descendants. Panel A shows a ventral view of the germband and panels B-F posterior views of the T2 limb. (G-L) Same renderings as in A-F with color masks showing in (G) the limb primordium, (H) the early limb bud, (I) the 2-partite limb with the first division between ischium/merus, (J) the 3-partite limb after the second division between basis/ischium, (K) the 6-partite limb after 3 more divisions between coxa/basis, propodus/dactylus and carpus/propodus, and (L) the final pattern made of the 7 segments after the carpus/merus division. Colored lines indicate the relationships between limb parts between consecutive stages. (M-R) Schematic representation of limb subdivision along the proximal-distal axis at the same time-points as in panels G-L. The rectangular lattice in panel M shows the 9 columns of cells at the 4-row-parasegment. White lines in panels N-R delineate the subdivisions of the T2 limb at the corresponding stages. The origin of each of the 7 segments in the differentiated T2 limb is shown with discs color-coded by segment.

In order to understand the cellular basis of the establishment of positional values along the proximal-distal axis during limb differentiation, we followed the fate of cells from the grid stage all the way to the fully differentiated T2 limb pattern (Figures 6A-6F). In particular, we tracked neighboring cells in column E4c (the anterior-dorsal cells E4c5-c8, not shown) and in column E5a (the posterior-dorsal cells E5a5-a8, shown in Figures 6A-6F) over 68 hours of development (84-151 hours AEL). These cells were ideal for reconstructing the proximal-distal axis at single-cell resolution because they mostly divided proximodistally forming elongated thin clones stretching from the coxa to the dactylus.

This lineage analysis demonstrated that the cells that gave rise to the proximal, medial and distal limb segments occupied distinct mediolateral positions in the germband grid at the 4-row-parasegment stage (Figure 6M) and distinct proximal-distal positions in the early limb bud (Figure 6N). Later on, analysis of the bipartite limb indicated that the proximal element gave rise to the proximal segments coxa, basis and ischium, while the distal element gave rise to the distal segments merus, carpus, propodus, and dactylus (Figure 6O). In other words, the first segmental subdivision occurred between ischium/merus and was followed by the basis/ischium subdivision (Figure 6P), the propodus/dactylus, carpus/propodus and coxa/basis subdivisions (Figure 6Q), and the carpus/merus subdivision (Figure 6R). Tracking analysis indicated that the cells forming the distal segments merus, carpus, propodus, and dactylus were not related by lineage, but by position as they originated from a medial territory around the intersection of the 4 compartments (Figure S9). During the subsequent limb elongation stages, these distal cells kept separate from more proximal cells at the prospective ischium/merus joint, suggesting that limb segments may represent secondary tissue subdivisions along the proximal-distal axis (Figure S9) (Milan and Cohen, 2000).

### Expression of limb patterning genes validates cellular models of *Parhyale* limb morphogenesis

To test the validity of our cellular models and make a first link between expression of limb patterning genes and morphogenetic cell behaviors, we cloned and analyzed by in situ hybridization the expression of the *Parhyale decapentaplegic (Ph-dpp)* gene that encodes a member of the Bone Morphogenetic Protein 2/4 class of signaling molecules. In *Drosophila* imaginal discs, Dpp signaling controls the dorsal cell fate in the leg, as well as growth via cell proliferation in the wing (Akiyama and Gibson, 2015; Brook and Cohen, 1996; Harmansa et al., 2015; Rogulja and Irvine, 2005; Svendsen et al., 2015). Therefore, probing *Ph-dpp* expression in forming *Parhyale* limb buds could provide a direct test for our cell-based predictions regarding the DV lineage restriction and the differential cell proliferation rates in the limb primordium.

Analysis of S18 embryos (96 hrs AEL) revealed alternating regions of high/moderate and low/no *Ph-dpp* expression in the anterior thoracic region (Figures 7A-7A’ and Figures S10A-S10A’). We used MaMuT to annotate with cellular resolution both the gene expression and the identity of cells in stained T2 limb buds. Acknowledging that the graded *Ph-dpp* expression at this stage obscured the precise limits of its expression, this analysis suggested that the region of high/moderate *Ph-dpp* expression was localized to rows E4c, E4d and E5a that mostly contribute to the presumptive dorsal compartment, while low/no *Ph-dpp* expression could be detected in the prospective ventral rows E4b anteriorly and E5b posteriorly (Figure 7A”). Furthermore, the *Ph-dpp* concentration gradient faded towards the medial (prospective ventral) columns and the border between high/moderate and low/no expressing cells was located in descendent cells from column 4 as also predicted by our *in silico* cellular analysis (Figures 7A-7A”). During the next 12 hrs (S19, 108 hrs AEL), the domain of strong *Ph-dpp* expression was more localized in the proximal-distal row of anterior-dorsal cells abutting the AP compartment boundary (Figures 7C-7C” and Figures S10C-S10C’).

**Figure 7.**
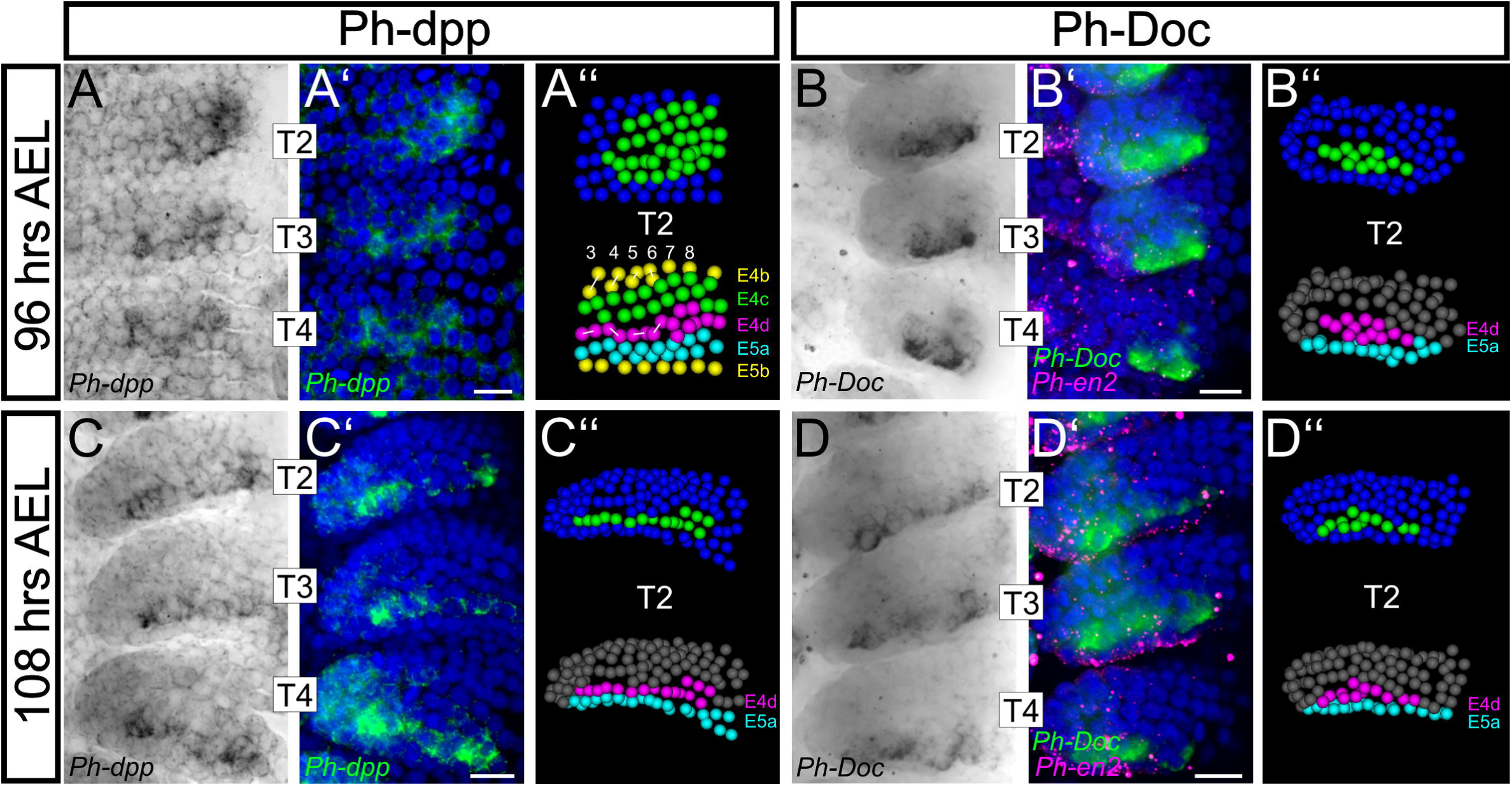
Analysis of Dpp expression and signaling corroborates cellular models of *Parhyale* limb development. (A-D) Brightfield images of T2, T3 and T4 limbs from S18 (top row, 96 hours After Egg Lay) and S19 (bottom row, 108 hours After Egg Lay) embryos stained by in situ hybridization for *Ph-dpp* (left column) and *Ph-Doc* (right column). (A’-D’) Same limbs as in panels A-D with the nuclear DAPI staining in blue overlaid with the *Ph-dpp* or *Ph-Doc* pattern false-colored in green. Embryos stained for *Ph-Doc* were co-hybridized with *Ph-en2* shown in magenta to label the posterior compartment. (A’’-D’’) MaMuT reconstructions of the T2 limbs shown in panels A-D’. The top panels are color-coded by gene expression with *Ph-dpp* or *Ph-Doc* expressing cells shown in green and non-expressing cells in blue. The bottom panels are color-coded by lineage with descendent cells from row a in cyan, row b in yellow, row c in green and row d in magenta. The column index number is shown at the top and white lines connect sister cells. All panels show ventral views with anterior to the top and ventral midline to the left. Scale bars are 20 μm.

To get an insight into the downstream effects of Dpp signaling in the *Parhyale* limb buds, we also analyzed by in situ hybridization the expression of the Tbx6/*Dorsocross* (*Doc*) gene that responds to high levels of Dpp signaling in the dorsal region of the *Drosophila* embryo and leg disc (Reim et al., 2003; Svendsen et al., 2015). Expression of the single *Doc* gene identified in *Parhyale* (*Ph-Doc*) was detected in a subset of the *Ph-dpp-* expressing cells in stage S18 (Figures 7B-7B’’ and Figures S10B-S10B’), while in stage S19 the two genes exhibited essentially identical strong expression in the anterior-dorsal cells abutting the AP boundary (Figures 7D-7D’’ and Figures S10D-S10D’). Importantly, in both stages analyzed, the anterior-dorsal limb cells experiencing high levels of *Ph-dpp* signaling and expressing *Ph-Doc* also exhibited the highest rates of cell proliferation (compare Figure 7 with Figure 4) providing strong correlative evidence for morphogen-dependent control of *Parhyale* limb growth.

Collectively, these results demonstrate how the reconstruction of cell lineages and behaviors can provide solid predictions and powerful contexts to study the expression and function of associated genes.

## DISCUSSION

In this article, we have established an integrated framework to study the cellular and genetic basis of developmental morphogenesis. By combining light-sheet microscopy with a newly developed versatile software for cell tracking and lineage reconstruction in large multi-view datasets, we have revealed the cellular architecture and cellular dynamics underlying organ morphogenesis in a non-conventional experimental model, and have provided new insights into the mechanisms underlying the three-dimensional epithelial remodeling during arthropod limb morphogenesis.

### MaMuT, a user-friendly tool for lineaging multi-dimensional image volumes

MaMuT’s unique feature that is currently not available in any other commercial or open-source software platforms for image data visualization and analysis is the capacity to track objects synergistically from all available views in multi-view LSFM recordings. This functionality has a number of very important advantages. Raw image stacks do not have to be fused into a single volume, which is computationally by far the most demanding step in multi-view LSFM image processing requiring access to high-end workstations and computer clusters or down-sampling of image volumes (Preibisch et al., 2014). The users also preserve the original redundancy of their data, which in many cases, including *Parhyale,* allows capturing each object of interest from two or more neighboring views that can be interpreted independently for a more accurate analysis. Finally, MaMuT allows users to analyze sub-optimal datasets that cannot be fused properly or may create fusion artifacts. Of course, combining the raw views with a high-quality fused volume is the best available option, especially when handling complex datasets with high cell densities.

While offering multi-view tracking, MaMuT delivers also a number of other important functionalities for the exploration of multi-dimensional datasets. First, MaMuT is a user-friendly turnkey software with a convenient interface for interactively navigating, inspecting and curating the image and annotation data. Any time-lapse series acquired by any microscopy modality that can be opened in Fiji can be also imported into MaMuT. Second, MaMuT offers a highly responsive and interactive navigation through multi-terabyte datasets fostered by the BigDataViewer architecture for image storage, loading and caching (Pietzsch et al., 2015). Individual z-stacks representing different views and/or channels of a multi-dimensional image dataset can be displayed independently (or in any combination) in multiple Viewer windows. Each Viewer can be also adjusted independently for color and brightness, zoom, translation and rotation of image stacks and browsing through time points. Third, MaMuT relies on TrackMate for the annotation model, which is a Fiji plugin developed for tracking purposes in conventional imaging datasets (Tinevez et al., 2016). Objects of interest like cells and nuclei (“spots”) can be selected synergistically from all available Viewers and followed over multiple time points to reconstruct their trajectories (“tracks”), division patterns and lineage information. Fourth, the created spots and tracks can be visualized and edited interactively in the Viewers and the TrackScheme lineage browser, and can be animated in the 3D Viewer window. For visual interpretation of the data, annotations can be colored based on the primary x, y, z, t and lineage information or derived numerical parameters such as velocity, displacement, division time and others. Fifth, lineages can be reconstructed in a fully manual, semi-automated or fully automated manner followed by manual curation if necessary. Thus, MaMuT can address various experimental designs ranging from analyzing a small subset of objects in the imaged volume to systems-wide analyses of all constituent parts. Sixth, all spot and track information is stored in an xml file that can be exported from MaMuT to other interfaces for more specialized analyses. Seventh, decentralized annotation by multiple users has been made possible by also developing a web service for remote access to large image volumes stored online. Last but not least, following on the tradition of the Fiji community for open-source distribution of biological image analysis software, MaMuT is provided freely and openly to the community, it is extensively documented and can be customized by other users.

### Multi-view LSFM is ideal to reveal the cellular basis of *Parhyale* morphogenesis

The LSFM imaging technology is empowering biologists to study developmental processes with unprecedented spatial and temporal resolution (Keller, 2013; Khairy and Keller, 2011; Schmied et al., 2014; Weber et al., 2014). We demonstrated here that in the multi-view mode it is particularly suited for imaging embryos of the crustacean amphipod *Parhyale hawaiensis* for several days. *Parhyale* has been already established as an attractive new model organism for developmental genetic and functional genomic studies supported by many experimental techniques, including transgenesis, CRISPR/Cas-mediated knock-out and knock-in approaches, genomic resources and other experimental tools (Kao et al., 2016; Martin et al., 2016; Pavlopoulos and Averof, 2005; Pavlopoulos et al., 2009; Stamataki and Pavlopoulos, 2016). By extending here the *Parhyale* toolkit with multi-view LSFM and the MaMuT reconstructions, it is now feasible to study gene expression and function in the context of single-cell resolution fate maps and cell pedigrees. Especially when it comes to appendage development, the *Parhyale* body plan provides exceptional biological material to probe the molecular and cellular basis of appendage patterning, growth and differentiation during embryogenesis and post-embryonic regeneration (Alwes et al., 2016; Browne et al., 2005; Konstantinides and Averof, 2014).

The tempo and mode of development has also important ramifications for *Parhyale* multi-view LSFM imaging and tracking. First, the slow tempo of development enabled to image the embryo at a very high spatial resolution through the acquisition of multiple and highly overlapping views without compromising the temporal resolution. Thus, *Parhyale* recordings can match or exceed the spatiotemporal resolution of other pioneering models in modern LSFM microscopy, like *Drosophila* and zebrafish, even when access to highest-speed instruments is not available. Second, due to the optical clarity of the embryo and positioning of the appendages on the surface of the developing embryo, all constituent cells were captured and tracked for systems-level quantitative analyses. Finally, the conserved, stereotypic and highly ordered organization of the post-naupliar ectoderm in *Parhyale* will allow to identify homologous cells and compare lineages, cell behaviors and associated genes between serially homologous structures in the same embryo, across embryos and even across malacostracan crustaceans.

### Cellular basis of arthropod limb morphogenesis: lessons from *Parhyale*

All our knowledge on the cellular basis of arthropod limb morphogenesis comes from studies in *Drosophila* (Fristrom, 1988; von Kalm et al., 1995). However, development of *Drosophila* limbs is extremely derived and not representative for many insects, much less arthropods in general. *Drosophila* limb specification, patterning, growth, and differentiation take place at distinct developmental stages during embryonic, larval and pupal development. On the contrary, all these processes come about during embryogenesis in most other arthropods, including *Parhyale,* and limbs are direct three-dimensional outgrowths of the embryonic body wall.

Using lineage-dependent or independent mechanisms, initially homogeneous fields of cells become progressively subdivided into distinct domains that differ in gene expression and often form and maintain sharp boundaries between themselves. Lineage-based subdivisions have been identified in the *Drosophila* wing and leg primordia during their compartmentalization along the AP axis in early embryogenesis, and during DV compartmentalization of the wing disc during early larval development (Garcia-Bellido et al., 1973; Steiner, 1976). These heritable subdivisions are based on the compartment-specific expression of selector genes and localized induction of signaling molecules (“organizers”) at the AP compartment boundaries in both legs and wings and the DV boundary of the wing (Dahmann et al., 2011; Garcia-Bellido, 1975; Lawrence and Struhl, 1996; Mann and Carroll, 2002). The mechanism underlying DV separation in the *Drosophila* leg disc is not clear yet. Classical lineaging experiments suggested a heritable DV division during larval development, although the clonal restriction was not entirely strict and was only detected in the anterior compartment but not the posterior one (supposedly due to its smaller size) (Steiner, 1976). It is also not clear yet whether the domains of expression of dorsal and ventral selector genes in the leg disc are coextensive with their corresponding compartments, and what are the cellular mechanisms contributing to regionalization of the leg along its DV axis (Brook and Cohen, 1996; Svendsen et al., 2015).

Beyond *Drosophila,* our understanding of the AP and DV organization in other arthropod limbs has relied so far entirely on gene expression studies. Expression of segment polarity genes, like *engrailed* and *wingless,* has demonstrated that the AP separation is widely conserved across the arthropods and takes place during the segmentation stages (Angelini and Kaufman, 2005; Patel et al., 1989). In *Parhyale,* the AP compartment boundary is established at the 1-row stage at the interface of neighboring parasegments (Browne et al., 2005; Scholtz et al., 1994). With the exception of descriptive gene expression studies (Janssen et al., 2008; Prpic et al., 2003), the mechanism, timing and position of the DV separation in arthropod limbs has remained unexplored at the cellular level. This limitation stems from the lack of methodologies to systematically generate and interpret the distribution and shape of mitotic recombination clones in non-model arthropods. Despite this limitation, we have been able to explore the cellular mechanisms underlying the developmental subdivisions in the *Parhyale* limb by analyzing the dynamics of digital clones in reconstructed T2 limbs. First, we confirmed that this approach was indeed successful in revealing the position and timing of the AP compartment boundary; the population of anterior cells (descendants from rows E4b, c and d) remained together and separate by a straight boundary from posterior cells (descendants from rows E5a and b). Second, by applying the same rules we revealed a putative heritable subdivision along the DV axis in two independently imaged and analyzed T2 limbs. The presumptive DV compartment boundary formed at the 4-row-parasegment stage between the E4b and c rows anteriorly, the E5a and b rows posteriorly, and between cells E4c4-c5, E4d3-d4 and E5a4-a5 medially. Third, expression of *Distal-less* (*Dll*), which is one of the earliest markers of limb specification, initiates at the 4-row-parasegment in the cells contributing to the appendage primordia (Browne et al., 2005; Hejnol and Scholtz, 2004). *Dll* expression is first detected in the d4 cell, which was one of the cells at the intersection of the AP and DV compartment boundaries. Fourth, this intersection coincided with the leading tip of the forming limb throughout epithelial remodeling and outgrowth.

These results demonstrate that the *Parhyale* limb perfectly conforms to the boundary model originally proposed by Hans Meihardt to whose memory this article is dedicated (Meinhardt, 1983). Meinhardt’s model postulated that a secondary developmental field, i.e. the proximal-distal axis of a limb that is specified during embryogenesis *de novo* relative to the main AP and DV body axes, initiates and is patterned around the intersection of the two AP and DV compartment boundaries.

The inference of the four constituent compartments provided a powerful framework to interpret the cell behaviors during the subsequent limb bud formation and elongation stages in a quantitative manner. This analysis strongly suggested that a combination of cellular (and associated molecular) mechanisms is at work to transform the two-dimensional embryonic epithelium into the three-dimensional outgrowing limb bud. First, there was a significant difference in cell proliferation rates between cells in the center (faster dividing) and the periphery (slower dividing) of the limb primordium from early primordium specification until the global elevation of cells above the level of the epithelium during limb bud formation. Such a “growth-based morphogenesis model” has been the dominant hypothesis for almost 50 years to explain the outgrowth of the vertebrate limb bud (Ede and Law, 1969; Hornbruch and Wolpert, 1970) - although it has been challenged recently (Boehm et al., 2010)) - but has never been implicated as the driving mechanism behind arthropod limb evagination. Second, the observed cell and tissue dynamics raised the question whether formation of the convex surface of the limb bud requires modulation of the basal adhesive properties of the dorsal and ventral cells, similar to the folding of the *Drosophila* wing bilayer (Fristrom et al., 1993). Third, the outgrowth of the limb was tightly associated - and presumably effected - by two patterned cell behaviors: i) an increased cell proliferation rate at the tip of the limb (at the intersection of the 4 compartments) resembling a putative growth zone which generates many of the new cells necessary for limb outgrowth, especially for the growing distal part of the limb; and ii) a strong bias in the orientation of mitotic divisions parallel to the proximal-distal axis of growth. Although cell proliferation rates were relatively heterogeneous in the rest of the limb, our analysis detected a second region of higher mitotic activity in the cells of the anterior compartment abutting the AP boundary both dorsally and ventrally. Fourth, the different proximal-distal domains of the *Parhyale* limb could be traced back to distinct mediolateral positions in the early germband stage. During limb bud formation and elongation, there was a transition and refinement of these positional values along the proximal-distal axis. Fifth, the temporal sequence of segmental subdivisions proceeded as follows (early < late): ischium/merus < basis/ischium < coxa/basis + propodus/dactylus + carpus/propodus < carpus/merus. Sixth, besides the early AP and DV lineage restrictions, we observed a non-heritable proximal-distal separation between cells in the ischium and the merus that were the first limb segments to separate, suggesting that limb segments may represent secondary tissue subdivisions.

Our systems-level approach demonstrates that the comprehensive imaging and fine-scale reconstruction of a developmental process can shed light into functionally interdependent patterning mechanisms operating across multiple scales. We expect that the availability of membrane fluorescent reporters for *Parhyale* live imaging will shed more light in the future on how cell rearrangements and cell shape changes contribute to limb morphogenesis.

### Reconciling genetic with cellular models of limb morphogenesis

As demonstrated in diverse developmental systems, compartment boundaries are important organizers of tissue patterning and growth through the expression of secreted morphogens and other signaling molecules (Dahmann et al., 2011; Lawrence and Struhl, 1996; Matsuda et al., 2016; Restrepo et al., 2014; Tabata and Takei, 2004). In the *Drosophila* leg disc, *engrailed* activates expression of the Hedgehog (Hh) secreted ligand in the posterior compartment, which in turn activates the Dpp and Wingless (Wg/Wnt-1) ligands at the AP boundary in the dorsal and ventral cells, respectively (Estella et al., 2012). Wg and Dpp create a concenration gradient with the highest level of the two ligands in the center of the disc and lower levels towards the periphery, and cooperate in the establishment of concentric domains of expression of a set of genes, referred to as limb gap genes, that pattern the limb proximal-distal axis: *Distal-less* distally, *dachshund* medially, *homothorax* and *extradenticle* proximally. Dpp and Wg signaling also act antagonistically to control the dorsal and ventral cell fates, respectively, through activation of the downstream selector T-box genes *optomotor blind/Dorsocross* dorsally and the *H15/midline* paralogs ventrally (Svendsen et al., 2015).

The proximal-distal expression of the limb gap genes is widely conserved in all arthropods studied, including *Parhyale* (Angelini and Kaufman, 2005; Browne et al., 2005; Prpic and Telford, 2008). Our analysis of *Ph-dpp* and *Ph-Doc* expression has also suggested a conserved function for Dpp signaling in dorsal fate specification and has provided extra independent support for a compartment-based mechanism to pattern the DV axis of *Parhyale* thoracic limbs. It has been argued that the stripy Wg and Dpp expression observed in the *Drosophila* leg disc is appropriate for patterning a twodimensional epithelium but not a three-dimensional outgrowing limb (Prpic et al., 2003). Expression at the tip of three-dimensional embryonic limbs has been demonstrated for Dpp in many studied insect and arthropod species (Angelini and Kaufman, 2005). This is not the case in *Parhyale,* where we detected a broad expression of *Ph-dpp* in the lateral (prospective dorsal) columns of the two-dimensional limb primordium. *Ph-wg* expression is not known in *Parhyale.* If it expressed in a complementary pattern to *Ph-dpp* in the medial (prospective ventral) columns, it could point to a similar logic for patterning the limb proximal-distal axis like in *Drosophila*. In fact, our reconstructions have suggested that the distal DV margin (experiencing the highest levels of Dpp and Wg signaling) is located between descendent cells from column 3, 4 and 5 (Figure 4). These are indeed the cells that contribute to the most distal limb segments (Figure 7).

Although the function of the Dpp morphogen gradient in patterning the *Drosophila* imaginal discs is well understood, its role in promoting growth is still controversial despite intensive efforts over the last decades (Akiyama and Gibson, 2015; Harmansa et al., 2015; Matsuda et al., 2016; Restrepo et al., 2014; Rogulja and Irvine, 2005). The dorsal-anterior cells expressing *Ph-dpp* and *Ph-Doc* were among the fastest dividing cells in the center of the limb primordium (compare Figure 4 with Figure 7). Later strong expression of *Ph-dpp* and *Ph-Doc* resolved into a proximal-distal row of cells of dorsal-anterior identity abutting the AP compratment boundary. Again, these cells displayed some of the highest proliferation rates quantified during limb outgrowth, suggesting a Dpp-dependent control of *Parhyale* limb growth. Thanks to the stereotyped and highly ordered *Parhyale* body plan, we anticipate that the LSFM imaging and tracking approaches described here, together with the recent application of CRSIPR/Cas-based methodologies for genome editing (Kao et al., 2016) will provide excellent material to further explore how morphogens like Dpp regulate form and function at cellular resolution.

## MATERIALS & METHODS

### Generation of transgenic *Parhyale* labeled with H2B-mRFPruby

*Parhyale* rearing, embryo collection, microinjection and generation of transgenic lines were carried out as previously described (Kontarakis and Pavlopoulos, 2014). To fluorescently label the chromatin in transgenic *Parhyale*, we fused the coding sequences of the *Drosophila melanogaster* histone *H2B* and the *mRFPruby* monomeric Red Fluorescent Protein and placed them under control of a strong *Parhyale* heat-inducible promoter (Pavlopoulos et al., 2009). *H2B* was amplified from genomic DNA with primers Dmel_H2B_F_NcoI (5’-TT AACCATGGCTCCGAAAACTAGTGGAAAG-3’) and Dmel_H2B_R_XhoI (5’-ACTTCTCGAGTTTAGAGCTGGTGTACTTGG-3’), and *mRFPruby* was amplified from plasmid pH2B-mRFPruby (Fischer et al., 2006) with primers mRFPruby_F_XhoI (5’-ACAACTCGAGATGGGCAAGCTTACC-3’) and mRFPruby_R_PspMOI (5’-TATTGGGCCCTTAGGATCCAGCGCCTGTGC-3’). The NcoI/XhoI-digested H2B and XhoI/PspOMI-digested mRFPruby fragments were cloned in a triple-fragment ligation into NcoI/NotI-digested vector pSL-PhHS-DsRed, placing *H2B-mRFPruby* under control of the *PhHS* promoter (Pavlopoulos et al., 2009). The *PhHS-H2B-mRFPruby-SV40polyA* cassette was then excised as an AscI fragment and cloned into the AscI-digested pMinos{3xP3-EGFP} vector (Pavlopoulos et al., 2004), generating plasmid pMi{3xP3-EGFP; PhHS-H2B-mRFPruby}. Three independent transgenic lines were established with this construct for heat-inducible expression of H2B-mRFPruby. The most strongly expressing line was selected for all applications. In this line, nuclear H2B-mRFPruby fluorescence plateaued about 12 hours after heat-shock and high levels of fluorescence persisted for at least 24 hours post heat-shock labeling chromatin in all cells throughout the cell cycle.

### Multi-view LSFM imaging of *Parhyale* embryos

To prepare embryos for LSFM imaging, 2.5-day old transgenic embryos (early germband stage S11; (Browne et al., 2005)) were heat-shocked for 1 hour at 37°C. About 12 hours later (stage S13), they were mounted individually in a cylinder of 1% low melting agarose (SeaPlaque, Lonza) inside a glass capillary (#701902, Brand GmbH) with their AP axis aligned parallel to the capillary. A 1:4000 dilution of red fluorescent beads (#F-Y050 microspheres, Estapor Merck) were included in the agarose as fiducial markers for multi-view reconstruction. During imaging, the embedded embryo was extruded from the capillary into the chamber filled with artificial seawater supplemented with antibiotics and antimycotics (FASWA; (Kontarakis and Pavlopoulos, 2014)). The FASWA in the chamber was replaced every 12 hours after each heat shock (see below). Embryos were imaged on a Zeiss Lightsheet Z.1 microscope equipped with a 20x/1.0 Plan Apochromat immersion detection objective (at 0.9 zoom) and two 10x/0.2 air illumination objectives producing two light-sheets 5.1 *μ*m thick at the waist and 10.2 *μ*m thick at the edges of a 488 *μ*m x 488 *μ*m field of view.

We started imaging *Parhyale* embryogenesis from 3 angles/views (the ventral side and the two ventral-lateral sides 45° apart from ventral view) during 3 to 4.5 days AEL to avoid damaging the dorsal thin extra-embryonic tissue, and continued imaging from 5 views (adding the two lateral sides 90° apart from ventral view) during 4.5 to 8 days AEL. A multi-view acquisition was made every 7.5 min at 26°C. The H2B-mRFPruby fluorescence levels were replenished regularly every 12 hours by raising the temperature in the chamber from 26°C to 37°C and heat-shocking the embryo for 1 hour. Each view (z-stack) was composed of 250 16-bit frames with voxel size 0.254 *μ*m x 0.254 *μ*m x 1 *μ*m. Each 1920 x 1920 pixel frame was acquired using two pivoting light-sheets to achieve a more homogeneous illumination and reduced image distortions caused by light scattering and absorption across the field of view. Each optical slice was acquired with a 561 nm laser and exposure time of 50 msec. With these conditions, *Parhyale* embryos were recorded routinely for a minimum of 4 days resulting on average in 192 time-points / 240K images / 1.7 TB of raw data per day.

### Multi-view LSFM image processing for 4D reconstruction of *Parhyale* embryogenesis

Image processing was carried out on a MS Windows 7 Professional 64-bit workstation with 2 Intel Xeon E5-2687W processors, 256 GB RAM (16 X DIMMs 16384 MB 1600 MHz ECC DDR3), 4.8 TB hard disk space (2 X 480 GB and 6 X 960 GB Crucial M500 SATA 6Gb/s SSD), 2 NVIDIA Quadro K4000 graphics cards (3 GB GDDR5). The workstation was connected through a 10 GB network interface to a MS Windows 2008 Server with 2 Intel Xeon E5-2680 processors, 196 GB RAM (24 X DIMM 8192 MB 1600 MHz ECC DDR3) and 144 TB hard disk space (36 X Seagate Constellation ES.3 4000 GB 7200 RPM 128 MB Cache SAS 6.0Gb/s). All major LSFM image data processing steps were done with software modules available through the Multiview Reconstruction Fiji plugin (http://imagej.net/Multiview-Reconstruction):

1) Preprocessing: Image data acquired on Zeiss Lightsheet Z.1 were saved as an array of czi files labeled with ascending indices, where each file represented one view (z-stack). czi files were first renamed into the “spim_TL{t}_Angle{a}.czi” filename, where t represented the time-point (e.g. 1 to 192 for a 1-day recording) and a the angle (e.g. 0 for left view, 45 for ventral-left view, 90 for ventral view, 135 for ventral-right view and 180 for right view), and then resaved as tif files (Schmied et al., 2016).
2) Bead-based spatial multi-view registration: In each time-point, each view was aligned to an arbitrary reference view fixed in 3D space (e.g. views 0, 45, 90, 135 aligned to 180) using the bead-based registration option (Preibisch et al., 2010). In each view, fluorescent beads scattered in the agarose were segmented with the Difference-of-Gaussian algorithm using a sigma value of 3 and an intensity threshold of 0.005. Corresponding beads were identified between views and were used to determine the affine transformation model that matched each view to the reference view within each time-point.
3) Fusion by multi-view deconvolution: Spatially registered views were down-sampled twice for time and memory efficient computations during the image fusion step. Input views were then fused into a single output 3D image with a more isotropic resolution using the Fiji plugin for multi-view deconvolution estimated from the point spread functions of the fluorescent beads (Preibisch et al., 2014). The same cropping area containing the entire imaged volume was selected for all time-points. In each time-point, the deconvolved fused image was calculated on GPU in blocks of 256x256x256 pixels with 7 iterations of the Efficient Bayesian method regularized with a Tikhonov parameter of 0.0006.
4) Bead-based temporal registration: To correct for small drifts of the embryo over the extended imaging periods (e.g. due to agarose instabilities), we stabilized the fused volume over time using the segmented beads (sigma = 1.8 and intensity threshold = 0.005) for temporal registration with the affine transformation model using an all-to-all matching within a sliding window of 5 time-points.
5) Computation of spatiotemporally registered fused volumes: Using the temporal registration parameters, we generated a stabilized time-series of the fused deconvolved 3D images.
6) 4D rendering: The *Parhyale* embryo was rendered over time from the spatiotemporally registered fused data using Fiji’s 3D Viewer.

### Lineage reconstruction with the Massive Multi-view Tracker (MaMuT)

MaMuT is a Fiji plugin that can be installed through the Fiji Updater. The source code for MaMuT is available on GitHub (https://github.com/fiji/MaMuT) and detailed tutorials and training datasets can be found at http://imagej.net/MaMuT. In brief, for lineaging purposes, the *Parhyale* multi-view LSFM raw z-stacks were registered spatiotemporally and the image data together with the registration parameters were converted into the custom HDF5/XML file formats utilized by the BigDataViewer and MaMuT plugins.

A new MaMuT annotation project was created for each dataset and appendage of interest. The nuclei contributing to the T2 limb were identified in the first time-point and tracked manually every 5 time-points except during mitosis, in which case we also tracked one time-point before and one after segregation of the daughter chromosomes during anaphase/telophase. To guarantee the accuracy of our lineage reconstructions, the position of each tracked nucleus was verified in at least two neighboring views and by slicing the data orthogonally in separate Viewer windows. All color-coded cell animations were created with MaMuT’s 3D Viewer, which is based on (Schmid et al., 2010).

The graph data structure in MaMuT can handle efficiently up to about a hundred thousand annotations. This number is well within the realm of manually generated annotations, but is normally exceeded by large-scale fully automated lineaging engines like TGMM. For this reason, we also provide users the option to crop the imported TGMM annotation in space and/or in time to make them compatible with MaMuT.

### In situ hybridization

In situ hybridizations were carried out as previously described (Rehm et al., 2009). The sequence accession number for *Parhyale decapentaplegic* (*Ph-dpp*) is XXX, for *Dorsocross* (*Ph-Doc*) XXX, and for *engrailed-2* (*Ph-en2*) XXX. Stained samples were imaged on a Zeiss 880 confocal microscope using the Plan-Apochromat 10x/0.45 and 20x/0.8 objectives. Images were adjusted for brightness/contrast and processed using Fiji (http://fiji.sc) and Photoshop (Adobe Systems Inc). For color overlays, the brightfield image of the *Ph-dpp* or *Ph-Doc* BCIP/NBT staining was inverted, false-colored green and merged with the fluorescent signal of the *Ph-en2* FastRed staining in magenta and the nuclear DAPI signal in blue.

## AUTHOR CONTRIBUTIONS

A.P. and C.W. conceived the project, A.P. and P.T. managed the project, A.P. and B.H. generated the *Parhyale* imaging data, J.Y.T. and T.P. developed MaMuT, T.P., S.P.,

P.J.K., P.T. and A.P. performed image analysis, E.S. carried out the in situ hybridizations, A.P. and C.W. analyzed the data and prepared the manuscript with input from all authors.

## ACKNOWLEDGEMENTS

We dedicate this article to the memory of Hans Meinhardt, whose insightful theoretical work advanced our understanding of developmental pattering mechanisms. We are really grateful to Michael Akam and Gerhard Scholtz for hosting and actively supporting the early phases of this project, Stephan Saalfeld for his help with image 3D rendering, Fernando Amat for his help with importing TGMM annotations into MaMuT, Angeliki-Myrto Farmaki for designing the MaMuT logo, and Michalis Averof for comments on the manuscript. C.W. was supported by the Einstein Foundation Berlin grant A-2012_114. J.Y.T. and S.S were supported by intramural funding from the Pasteur Institute. T.P. and P.T. were supported by the core funding from Max Planck Society and the European Research Council Community's Seventh Framework Program, grant agreement 260746. A.P., E.S. and P.J.K. were supported by the Howard Hughes Medical Institute. A.P. was also supported by a Marie Curie Intra-European fellowship.

## SUPPLEMENTAL INFORMATION

**Figure S1. MaMuT layout**

(A-C) The three tabs of the MaMuT control panel. (A) The Views tab is used to launch and control the different displays of the image data and annotations. (B) The Annotation tab is used to define the temporal sampling during manual lineaging and the parameters for semi-automated tracking. (C) The Actions tab allows users to generate movies of tracked objects of interest or merge independent MaMuT annotations of the same image dataset into a single file. (D) The visibility and grouping panel allows users to organize views into groups and display them overlaid in the same Viewer window. (E) The brightness and color panel is used to adjust brightness, contrast and color of the image data in the Viewer windows. For example, the three different views are shown here in cyan, blue and yellow, respectively. (F) The MaMuT help menu with the default mouse and keyboard operations. The default key bindings can be modified by the user. (G-L) The MaMuT Viewer windows display the raw image data. The user interacts with the data to create and edit annotations through these Viewers and the TrackScheme window. The user can open as many Viewer windows as required for accurate tracking and can display the image data in any useful scale and orientation (here two orientations per view). (M) The TrackScheme window is the dedicated lineage browser and editor where tracks are arranged from left to right and time-points from top to bottom. (N) The 3D Viewer window shows animations of the tracked objects as spheres without the image data. The TrackScheme, the 3D Viewer and all open Viewer windows are synced (with the current selection highlighted in bright green) and annotations can be color-coded according to various parameters extracted from the data.

**Figure S2. Lineage reconstruction of anterior-posterior rows in the *Parhyale* thoracic limb primordium**

(A-E) Lateral views of a *Parhyale* embryo reconstructed from a multi-view LSFM acquisition at the indicated developmental stages shown in hours (h) after egg-lay (AEL). The yellow mask indicates the second thoracic appendage (T2) on the left side that was reconstructed at a single-cell resolution. (F-J’) Tracked cells contributing to the outgrowing T2 limb belong to the E4 and E5 parasegments, and are color-coded by the parasegmental row they belong to: a row in cyan, b rows in yellow, c row in green and d row in magenta. (F) Ventral view of the limb primordium at the 4-row-parasegment stage at 84 h AEL. Horizontal lines separate rows a to d along the AP axis. (G) Ventral view of the limb primordium during early eversion at 96 h AEL. (H-J) Dorsal views of (H) limb bud at 103 h AEL, (I) initial limb elongation at 114 h AEL and (J) later elongation phase at 123 h AEL. (H’-J’) Ventral views, rotated 180° relative to H-J. Note the absence of cell mixing at the AP compartment boundary between the anterior E4c/E4d cells (green/magenta) and the posterior E5a cells (cyan), as well as at the DV compartment boundary between the ventral E4b/E5b cells (yellow) and their dorsal anterior E4c neighbors (green) and dorsal posterior E5a neighbors (cyan).

**Figure S3. Lineage reconstruction of dorsal-ventral columns in the *Parhyale* thoracic limb primordium**

(A-E) Lateral views of a *Parhyale* embryo reconstructed from a multi-view LSFM acquisition at the indicated developmental stages shown in hours (h) after egg-lay (AEL). The yellow mask indicates the second thoracic appendage (T2) on the left side that was reconstructed at a single-cell resolution. (F-J’) Tracked cells contributing to the outgrowing T2 limb are color-coded by the dorsal-ventral column they belong to: column 3 in yellow, 4 in orange, 5 in red, 6 in green, 7 in cyan, 8 in blue, and 9 in magenta. (F) Ventral view of the limb primordium at the 4-row-parasegment stage at 84 h AEL. Vertical lines separate columns along the DV axis. (G) Ventral view of the limb primordium during early eversion at 96 h AEL. (H-J) Dorsal views of (H) limb bud at 103 h AEL, (I) initial limb elongation at 114 h AEL and (J) later elongation phase at 123 h AEL. (H’-J’) Ventral views, rotated 180° relative to H-J. Note the cell mixing and irregular boundaries between descendants from neighboring columns.

**Figure S4. Cell lineage tree of the *Parhyale* T2 limb**

Each track resembles one or two of the 34 constituent cells of the T2 limb primordium color-coded by their compartmental identity: anterior-dorsal in dark green, anterior-ventral in dark magenta, posterior-dorsal in light green, and posterior-ventral in light magenta. Tracks labeled with the names of the corresponding cells are arranged horizontally and time-points are arranged vertically.

**Figure S5. Confirmation of early limb compartmentalization in another imaged and analyzed dataset**

(A-D) Lateral views of a *Parhyale* embryo reconstructed from a multi-view acquisition on a Zeiss LSFM prototype instrument that offered one-sided illumination with single-sided detection. One side of this embryo was imaged from 3 views 40° apart (ventral, ventral-left and left) every 7.5 min. These views were registered using fluorescent beads and were fused by multi-view deconvolution at the indicated developmental stages shown in hours (h) after egg-lay (AEL). The yellow mask indicates the second thoracic appendage (T2) that was reconstructed at a single-cell resolution. (F-I’) Tracked cells contributing to the outgrowing T2 limb color-coded by their compartmental identity: anterior-dorsal (AD; dark green), anterior-ventral (AV; dark magenta), posterior-dorsal (PD; light green), and posterior-ventral (PV; light magenta). (F) Ventral view of the limb primordium at the 4-row-parasegment stage at 86 h AEL. Horizontal lines separate rows a to d along the AP axis, and vertical lines separate columns 3 to 9 along the DV axis. (F’) Posterior view, rotated 90° relative to F. (G) Ventral view of the limb primordium during eversion at 94 h AEL. (G’) Posterior view, rotated 90° relative to G. The cells close to the intersection of the four compartments (yellow arrow) have risen above the level of the epithelium. (H-I) Dorsal views of (H) limb bud at 98 h AEL and (I) initial limb elongation at 101 h AEL. (H’-I’) Ventral views, rotated 180° relative to H-I. The intersection of the AP and DV compartment boundaries (yellow arrows) marks the tip of the limb. Cell tracking and lineage reconstructions shown in panels F-I’ were carried out with the SIMI°BioCell software (Schnabel et al., 1997) on a cropped single view of this dataset. Despite this limitation, we found exactly the same lineage restrictions as in the more complete reconstructions: cells in the anterior compartment (E4b, c and d rows) remained together and separate by a straight boundary from the cells in the posterior compartment (E5a and b rows). Likewise, no cell mixing was detected across the dorsal-ventral compartment boundary that extended again between the E4b and c rows anteriorly, between cells E4c4-c5, E4d3-d4 and E5a4-a5 distally, and between the E5a and b rows posteriorly.

**Figure S6. Cell intercalation during outgrowth of the *Parhyale* thoracic limb**

(A-D) Intercalation of descendant cells from the E4d6 lineage (cyan) between cells from the E4c6 lineage (yellow). (E-H) Intercalation of descendant cells from the E4d3 lineage (yellow) between cells from the E4c4 lineage (cyan). The limb has been reconstructed at the indicated developmental stages shown in hours (h) after egg-lay (AEL) and the constituent cells have been color-coded by their compartmental identity: anterior-dorsal (AD) in dark green, anterior-ventral (AV) in dark magenta, posterior-dorsal (PD) in light green, and posterior-ventral (PV) in light magenta.

**Figure S7. Digital clonal analysis in the *Parhyale* thoracic limb**

Digital clones (shown in bright green) for each one of the 34 constituent cells of the T2 limb primordium visualized at 114 hours (h) after egg-lay (AEL). The name of each cell and its position in the grid at the 4-row-parasegment (at 84 h AEL) is shown in each panel in the top left and bottom left corner, respectively. Besides the cells of the clone, the rest cells have been color-coded by their compartmental identity: anterior-dorsal (AD) in dark green, anterior-ventral (AV) in dark magenta, posterior-dorsal (PD) in light green, and posterior-ventral (PV) in light magenta.

**Figure S8. Alternative quantifications of cell proliferation rates in the *Parhyale* thoracic limb**

(A-E) Tracked cells making up the T2 limb shown at the indicated hours (h) after egg-lay (AEL) and color-coded by their compartmental identity: anterior-dorsal (AD) in dark green, anterior-ventral (AV) in dark magenta, posterior-dorsal (PD) in light green, and posterior-ventral (PV) in light magenta. (F-J) Same stages as in A-E color-coded by the average cell cycle length of each cell according to the scale shown on the right. (F’-J’) Same stages as in A-E color-coded by the absolute cell cycle length of each cell according to the scale shown on the right. (F’’-J’’) Same stages as in A-E color-coded by the average cell cycle length of each track according to the scale shown on the right. The AP and DV compartment boundaries are indicated by the cyan and green line, respectively. During the early stages, all analyses-irrespective of the method of quantification - demonstrate that a group of central cells in the limb primordium divide faster compared to peripheral cells. During later stages, the higher cell proliferation rates at the tip of the limb and at the AP compartment boundary is more pronounced in the calculation of the average cell cycle length of each cell. Gray cells indicate cells for which measurements are not applicable.

**Figure S9. Proximal-distal lineage separation in the growing *Parhyale* thoracic limb**

(A-E) Tracked cells contributing to the outgrowing T2 limb color-coded by their compartmental identity: anterior-dorsal (AD; dark green), anterior-ventral (AV; dark magenta), posterior-dorsal (PD; light green), and posterior-ventral (PV; light magenta). (A) Ventral view of the limb primordium at the 4-row-parasegment stage at 84 hours (h) after egg-lay (AEL). (B) Ventral view of the limb primordium during early eversion at 96 h AEL. (C) Dorsal view of limb bud at 103 h AEL. Posterior views of (D) initial limb elongation at 114 h AEL and (J) later elongation phase at 123 h AEL. (F-J) Same stages and views as in A-E with cells contributing to the proximal (p) leg segments (coxa, basis, and ischium) shown in cyan and cells contributing to the distal (d) leg segments (merus, carpus, propodus, and dactylus) shown in yellow. Progenitor cells giving rise to both proximal and distal leg segments are shown in bright green. (K-M) Later stages of limb segmentation at (K) 132 h AEL, (L) 140 h AEL and (M) 150 h AEL. In these panels, the T2 limb has been rendered in posterior view and superimposed with the tracked cells (descendant cells from posterior-dorsal progenitors E5a5-a8 shown as dots) stretching along the limb proximal-distal axis. Note that the proximal cells (in cyan) and the distal cells (in yellow) stop mixing at the ischium/merus joint (demarcated with the white line in K-M) after about 110 h AEL. (N) Cell lineage tree of the *Parhyale* T2 limb where the tracks have been color-coded by their proximal-distal identity: proximal identity in cyan, distal identity in yellow, and mixed identity in green. Proximal and distal cells are not related by lineage, but by position as they originate from the peripheral and medial territories of the limb primordium, respectively.

**Figure S10. Expression of *Ph-dpp* and *Ph-Doc* during *Parhyale* limb bud formation**

(A-D) Brightfield images of S18 (top row, 96 hours After Egg Lay) and S19 (bottom row, 108 hours After Egg Lay) embryos stained by in situ hybridization for *Ph-dpp* (left column) and *Ph-Doc* (right column). (A’-D’) Same embryos as in panels A-D with the nuclear DAPI staining in blue overlaid with the *Ph-dpp* or *Ph-Doc* pattern false-colored in green. Embryos stained for *Ph-Doc* were co-hybridized with *Ph-en2* shown in magenta to label the posterior compartment. All panels show ventral views with anterior to the top. Rectangles indicate the T2, T3 and T4 limbs shown in Figure 7. Scale bars are 100 μm.

## REFERENCES

Akiyama, T., and Gibson, M.C. (2015). Decapentaplegic and growth control in the developing Drosophila wing. Nature 527, 375–378.

Alwes, F., Enjolras, C., and Averof, M. (2016). Live imaging reveals the progenitors and cell dynamics of limb regeneration. Elife 5.

Amat, F., Hockendorf, B., Wan, Y., Lemon, W.C., McDole, K., and Keller, P.J. (2015). Efficient processing and analysis of large-scale light-sheet microscopy data. Nat Protoc 10, 1679–1696.

Amat, F., Lemon, W., Mossing, D.P., McDole, K., Wan, Y., Branson, K., Myers, E.W., and Keller, P.J. (2014). Fast, accurate reconstruction of cell lineages from large-scale fluorescence microscopy data. Nat Methods 11, 951–958.

Angelini, D.R., and Kaufman, T.C. (2005). Insect appendages and comparative ontogenetics. Dev Biol 286, 57–77.

Baena-Lopez, L.A., Baonza, A., and Garcia-Bellido, A. (2005). The orientation of cell divisions determines the shape of Drosophila organs. Curr Biol 15, 1640–1644.

Boehm, B., Westerberg, H., Lesnicar-Pucko, G., Raja, S., Rautschka, M., Cotterell, J., Swoger, J., and Sharpe, J. (2010). The role of spatially controlled cell proliferation in limb bud morphogenesis. PLoS Biol 8, e1000420.

Brook, W.J., and Cohen, S.M. (1996). Antagonistic interactions between wingless and decapentaplegic responsible for dorsal-ventral pattern in the Drosophila Leg. Science 273, 1373–1377.

Browne, W.E., Price, A.L., Gerberding, M., and Patel, N.H. (2005). Stages of embryonic development in the amphipod crustacean, Parhyale hawaiensis. Genesis 42, 124–149.

Buckingham, M.E., and Meilhac, S.M. (2011). Tracing cells for tracking cell lineage and clonal behavior. Dev Cell 21, 394–409.

Chhetri, R.K., Amat, F., Wan, Y., Hockendorf, B., Lemon, W.C., and Keller, P.J. (2015). Whole-animal functional and developmental imaging with isotropic spatial resolution. Nat Methods 12, 1171–1178.

Dahmann, C., Oates, A.C., and Brand, M. (2011). Boundary formation and maintenance in tissue development. Nat Rev Genet 12, 43–55.

Dohle, W., Gerberding, M., Hejnol, A., and Scholtz, G. (2004). Cell lineage, segment differentiation, and gene expression in crustaceans. In Evolutionary Developmental Biology of Crustacea, G. Scholtz, ed. (Lisse, The Netherlands: A. A. Balkema Publishers).

Ede, D.A., and Law, J.T. (1969). Computer simulation of vertebrate limb morphogenesis. Nature 221, 244–248.

Estella, C., Voutev, R., and Mann, R.S. (2012). A dynamic network of morphogens and transcription factors patterns the fly leg. Curr Top Dev Biol 98, 173–198.

Fischer, M., Haase, I., Wiesner, S., and Muller-Taubenberger, A. (2006). Visualizing cytoskeleton dynamics in mammalian cells using a humanized variant of monomeric red fluorescent protein. FEBS Lett 580, 2495–2502.

Fristrom, D. (1988). The cellular basis of epithelial morphogenesis. A review. Tissue Cell 20, 645–690.

Fristrom, D., Wilcox, M., and Fristrom, J. (1993). The distribution of PS integrins, laminin A and F-actin during key stages in Drosophila wing development. Development 117, 509–523.

Garcia-Bellido, A. (1975). Genetic control of wing disc development in Drosophila. Ciba Found Symp 0, 161–182.

Garcia-Bellido, A., Ripoll, P., and Morata, G. (1973). Developmental compartmentalisation of the wing disk of Drosophila. Nat New Biol 245, 251–253.

Hannibal, R.L., Price, A.L., and Patel, N.H. (2012). The functional relationship between ectodermal and mesodermal segmentation in the crustacean, Parhyale hawaiensis. Dev Biol 361, 427–438.

Harmansa, S., Hamaratoglu, F., Affolter, M., and Caussinus, E. (2015). Dpp spreading is required for medial but not for lateral wing disc growth. Nature 527, 317–322.

Hejnol, A., and Scholtz, G. (2004). Clonal analysis of Distal-less and engrailed expression patterns during early morphogenesis of uniramous and biramous crustacean limbs. Dev Genes Evol 214, 473–485.

Hornbruch, A., and Wolpert, L. (1970). Cell division in the early growth and morphogenesis of the chick limb. Nature 226, 764–766.

Huisken, J., Swoger, J., Del Bene, F., Wittbrodt, J., and Stelzer, E.H. (2004). Optical sectioning deep inside live embryos by selective plane illumination microscopy. Science 305, 1007–1009.

Janssen, R., Feitosa, N.M., Damen, W.G., and Prpic, N.M. (2008). The T-box genes H15 and optomotor-blind in the spiders Cupiennius salei, Tegenaria atrica and Achaearanea tepidariorum and the dorsoventral axis of arthropod appendages. Evol Dev 10, 143–154.

Kao, D., Lai, A.G., Stamataki, E., Rosic, S., Konstantinides, N., Jarvis, E., Di Donfrancesco, A., Pouchkina-Stancheva, N., Semon, M., Grillo, M., et al. (2016). The genome of the crustacean Parhyale hawaiensis, a model for animal development, regeneration, immunity and lignocellulose digestion. Elife 5.

Keller, P.J. (2013). Imaging morphogenesis: technological advances and biological insights. Science 340, 1234168.

Keller, P.J., Schmidt, A.D., Wittbrodt, J., and Stelzer, E.H. (2008). Reconstruction of zebrafish early embryonic development by scanned light sheet microscopy. Science 322, 1065–1069.

Khairy, K., and Keller, P.J. (2011). Reconstructing embryonic development. Genesis 49, 488–513.

Konstantinides, N., and Averof, M. (2014). A common cellular basis for muscle regeneration in arthropods and vertebrates. Science 343, 788–791.

Kontarakis, Z., and Pavlopoulos, A. (2014). Transgenesis in non-model organisms: the case of Parhyale. Methods Mol Biol 1196, 145–181.

Krzic, U., Gunther, S., Saunders, T.E., Streichan, S.J., and Hufnagel, L. (2012). Multiview light-sheet microscope for rapid in toto imaging. Nat Methods 9, 730–733.

Lawrence, P.A., and Struhl, G. (1996). Morphogens, compartments, and pattern: lessons from drosophila? Cell 85, 951–961.

Lecuit, T., and Le Goff, L. (2007). Orchestrating size and shape during morphogenesis. Nature 450, 189–192.

Liu, Z., and Keller, P.J. (2016). Emerging Imaging and Genomic Tools for Developmental Systems Biology. Dev Cell 36, 597–610.

Mann, R.S., and Carroll, S.B. (2002). Molecular mechanisms of selector gene function and evolution. Curr Opin Genet Dev 12, 592–600.

Martin, A., Serano, J.M., Jarvis, E., Bruce, H.S., Wang, J., Ray, S., Barker, C.A., O'Connell, L.C., and Patel, N.H. (2016). CRISPR/Cas9 Mutagenesis Reveals Versatile Roles of Hox Genes in Crustacean Limb Specification and Evolution. Curr Biol 26, 14–26.

Matsuda, S., Harmansa, S., and Affolter, M. (2016). BMP morphogen gradients in flies. Cytokine Growth Factor Rev 27, 119–127.

Meinhardt, H. (1983). Cell determination boundaries as organizing regions for secondary embryonic fields. Dev Biol 96, 375–385.

Milan, M., and Cohen, S.M. (2000). Subdividing cell populations in the developing limbs of Drosophila: do wing veins and leg segments define units of growth control? Dev Biol 217, 1–9.

Patel, N.H., Kornberg, T.B., and Goodman, C.S. (1989). Expression of engrailed during segmentation in grasshopper and crayfish. Development 107, 201–212.

Pavlopoulos, A., and Averof, M. (2005). Establishing genetic transformation for comparative developmental studies in the crustacean Parhyale hawaiensis. Proc Natl Acad Sci U S A 102, 7888–7893.

Pavlopoulos, A., Berghammer, A.J., Averof, M., and Klingler, M. (2004). Efficient transformation of the beetle Tribolium castaneum using the Minos transposable element: quantitative and qualitative analysis of genomic integration events. Genetics 167, 737–746.

Pavlopoulos, A., Kontarakis, Z., Liubicich, D.M., Serano, J.M., Akam, M., Patel, N.H., and Averof, M. (2009). Probing the evolution of appendage specialization by Hox gene misexpression in an emerging model crustacean. Proc Natl Acad Sci U S A 106, 13897–13902.

Pietzsch, T., Saalfeld, S., Preibisch, S., and Tomancak, P. (2015). BigDataViewer: visualization and processing for large image data sets. Nat Methods 12, 481–483.

Preibisch, S., Amat, F., Stamataki, E., Sarov, M., Singer, R.H., Myers, E., and Tomancak, P. (2014). Efficient Bayesian-based multiview deconvolution. Nat Methods 11, 645–648.

Preibisch, S., Saalfeld, S., Schindelin, J., and Tomancak, P. (2010). Software for bead-based registration of selective plane illumination microscopy data. Nat Methods 7, 418–419.

Prpic, N.M., Janssen, R., Wigand, B., Klingler, M., and Damen, W.G. (2003). Gene expression in spider appendages reveals reversal of exd/hth spatial specificity, altered leg gap gene dynamics, and suggests divergent distal morphogen signaling. Dev Biol 264, 119–140.

Prpic, N.M., and Telford, M.J. (2008). Expression of homothorax and extradenticle mRNA in the legs of the crustacean Parhyale hawaiensis: evidence for a reversal of gene expression regulation in the pancrustacean lineage. Dev Genes Evol 218, 333–339.

Rauskolb, C. (2001). The establishment of segmentation in the Drosophila leg. Development 128, 4511–4521.

Rehm, E.J., Hannibal, R.L., Chaw, R.C., Vargas-Vila, M.A., and Patel, N.H. (2009). In situ hybridization of labeled RNA probes to fixed Parhyale hawaiensis embryos. Cold Spring Harb Protoc 2009, pdb prot5130.

Reim, I., Lee, H.H., and Frasch, M. (2003). The T-box-encoding Dorsocross genes function in amnioserosa development and the patterning of the dorsolateral germ band downstream of Dpp. Development 130, 3187–3204.

Restrepo, S., Zartman, J.J., and Basler, K. (2014). Coordination of patterning and growth by the morphogen DPP. Curr Biol 24, R245–255.

Rogulja, D., and Irvine, K.D. (2005). Regulation of cell proliferation by a morphogen gradient. Cell 123, 449–461.

Schindelin, J., Arganda-Carreras, I., Frise, E., Kaynig, V., Longair, M., Pietzsch, T., Preibisch, S., Rueden, C., Saalfeld, S., Schmid, B., et al. (2012). Fiji: an open-source platform for biological-image analysis. Nat Methods 9, 676–682.

Schmid, B., Schindelin, J., Cardona, A., Longair, M., and Heisenberg, M. (2010). A high-level 3D visualization API for Java and ImageJ. BMC Bioinformatics 11, 274.

Schmied, C., Stamataki, E., and Tomancak, P. (2014). Open-source solutions for SPIMage processing. Methods Cell Biol 123, 505–529.

Schmied, C., Steinbach, P., Pietzsch, T., Preibisch, S., and Tomancak, P. (2016). An automated workflow for parallel processing of large multiview SPIM recordings. Bioinformatics 32, 1112–1114.

Schnabel, R., Hutter, H., Moerman, D., and Schnabel, H. (1997). Assessing normal embryogenesis in Caenorhabditis elegans using a 4D microscope: variability of development and regional specification. Dev Biol 184, 234–265.

Scholtz, G., Patel, N.H., and Dohle, W. (1994). Serially homologous engrailed stripes are generated via different cell lineages in the germ band of amphipod crustaceans (Malacostraca, Peracarida). Int J Dev Biol 38, 471–478.

Stamataki, E., and Pavlopoulos, A. (2016). Non-insect crustacean models in developmental genetics including an encomium to Parhyale hawaiensis. Curr Opin Genet Dev 39, 149–156.

Steiner, E. (1976). Establishment of compartments in the developing leg imaginal discs of Drosophila melanogaster. Wilhelm Roux Arch Dev Biol 180, 9–30.

Svendsen, P.C., Ryu, J.R., and Brook, W.J. (2015). The expression of the T-box selector gene midline in the leg imaginal disc is controlled by both transcriptional regulation and cell lineage. Biol Open 4, 1707–1714.

Swoger, J., Verveer, P., Greger, K., Huisken, J., and Stelzer, E.H. (2007). Multi-view image fusion improves resolution in three-dimensional microscopy. Opt Express 15, 8029–8042.

Tabata, T., and Takei, Y. (2004). Morphogens, their identification and regulation. Development 131, 703–712.

Tinevez, J.Y., Perry, N., Schindelin, J., Hoopes, G.M., Reynolds, G.D., Laplantine, E., Bednarek, S.Y., Shorte, S.L., and Eliceiri, K.W. (2016). TrackMate: an open and extensible platform for single-particle tracking. Methods.

Tomer, R., Khairy, K., Amat, F., and Keller, P.J. (2012). Quantitative high-speed imaging of entire developing embryos with simultaneous multiview light-sheet microscopy. Nat Methods 9, 755–763.

Truong, T.V., Supatto, W., Koos, D.S., Choi, J.M., and Fraser, S.E. (2011). Deep and fast live imaging with two-photon scanned light-sheet microscopy. Nat Methods 8, 757–760.

von Kalm, L., Fristrom, D., and Fristrom, J. (1995). The making of a fly leg: a model for epithelial morphogenesis. Bioessays 17, 693–702.

Weber, M., Mickoleit, M., and Huisken, J. (2014). Light sheet microscopy. Methods Cell Biol 123, 193–215.

Wolff, C., and Scholtz, G. (2008). The clonal composition of biramous and uniramous arthropod limbs. Proc Biol Sci 275, 1023–1028.

Wu, Y., Wawrzusin, P., Senseney, J., Fischer, R.S., Christensen, R., Santella, A., York, A.G., Winter, P.W., Waterman, C.M., Bao, Z., et al. (2013). Spatially isotropic fourdimensional imaging with dual-view plane illumination microscopy. Nat Biotechnol 31, 1032–1038.

